# Mpl is activated by dimers of MPN-linked calreticulin mutants stabilized by disulfide bonds and ionic interactions

**DOI:** 10.1101/2020.09.13.295485

**Authors:** Arunkumar Venkatesan, Jie Geng, Malathi Kandarpa, Sanjeeva Joseph Wijeyesakere, Ashwini Bhide, Moshe Talpaz, Irina D. Pogozheva, Malini Raghavan

## Abstract

Myeloproliferative neoplasms (MPNs) are frequently driven by insertions and deletions within the gene encoding calreticulin (CRT). CRT_Del52_ and CRT_Ins5_ are recurrent mutations. Although oncogenic transformation requires both mutated CRT and the myeloproliferative leukemia protein (Mpl), the molecular mechanism of CRT-mediated constitutive activation of Mpl is unknown. Our studies reveal that the novel C-domain of CRT_Del52_ encodes specificity both for Mpl binding and for disulfide-mediated CRT dimerization. Disulfide-stabilized CRT_Del52_ dimers and multimers are observed in MPN patient-derived platelet lysates and in transfected mammalian cells. Cysteine mutations within both the novel C-domain (C400A and C404A) and the conserved N-domain (C163A) of CRT_Del52_ are required to reduce disulfide-mediated dimers and multimers of CRT_Del52_. Based on these data and published structures of crystalized CRT oligomers, we tested the relevance of ionic interactions between charged residues proximal to C163 at the N-domain dimerization interface. Charge alteration at these residues affected dimerization and multimerization of both wild type and CRT_Del52_. Elimination of intermolecular disulfides and disruption of ionic interactions at both proposed dimerization interfaces was required to abrogate the ability of CRT_Del52_ to induce cytokine-independent cell proliferation via Mpl. Based on these findings, we propose a structural model of the Mpl-activating CRT_Del52_ unit as a covalently-linked dimer that is stabilized by disulfides and ionic interactions at both the C-domain and N-domain. MPNs exploit a natural dimerization interface of CRT combined with C-domain gain-of-functions to achieve cell transformation.

## Introduction

Myeloproliferative neoplasms (MPNs) comprising Polycythemia Vera (PV), Essential Thrombocythemia (ET) and Primary Myelofibrosis (MF) are hematopoietic stem cell disorders characterized by the overproduction of myeloid lineage cells (reviewed in reference (1)). Somatic mutations resulting from deletions or insertions in exon 9 of the *CALR* gene were identified in the majority of patients with PV, ET, and MF who were negative for mutations in Janus Kinase 2 (JAK2) and in the thrombopoietin receptor/myeloproliferative leukemia protein (Mpl) (2, 3). A majority of the *CALR*-mutated patients have one of two gene variants: type 1 with a 52 bp deletion (Del52) or type 2 with a 5 bp insertion (Ins5) (2, 3). The mutations change the sequence of the acidic C-terminus of CRT to a basic sequence, and cause loss of the endoplasmic reticulum (ER) retention KDEL sequence (2, 3).

Calreticulin (CRT) is an ER calcium-binding chaperone that functions in the folding and assembly of glycoproteins (4, 5). CRT contains three domains, a lectin-like N-terminal domain (N-domain) containing the glycan and high-affinity calcium binding sites (6, 7), an elongated hairpin-like P domain containing the co-chaperone-binding site (8) and a mainly α-helical acidic C-terminal domain (C-domain), which contains multiple low-affinity/Ca^2+^ binding sites (9–11). CRT has specificity for monoglucosylated N-glycans on substrate proteins, which are transiently acquired during glycoprotein maturation in the ER.

MPN-linked CRT mutants induce specific amplification of the megakaryocyte lineage of cells and increase platelet production (12–16). CRT mutants mediate constitutive activation of Mpl and downstream signaling pathways (12–15, 17–20). Studies of the direct interaction between Mpl and MPN-linked CRT mutants by co-immunoprecipitation analyses demonstrated a critical role of glycan-binding site residues of CRT (13, 20, 21) in the recognition of sugars linked to N117 of Mpl (13, 22). Since both wild type and mutant CRT are, in principle, capable of glycan-mediated interactions with Mpl, glycan-binding alone cannot account for mutant CRT-mediated Mpl activation. The novel basic amino acids in the C-termini of mutant CRTs were shown to be also critical for inducing cell proliferation and developing the ET phenotype (15, 18, 19). Studies of truncated mutants revealed that residues 376-383 from the C-terminus of CRT_Del52_ are required to activate Mpl-mediated signaling (20). However, the molecular contacts between Mpl and the C-terminal tail of CRT mutants remain undefined. Here we demonstrate a direct role of the C-terminal tail of CRT_Del52_ in Mpl binding and in conferring Mpl specificity to CRT_Del52_.

Ligand-induced dimerization of receptor molecules is an established paradigm for signal transduction mediated by cytokine receptors (23, 24) including the erythropoietin (Epo) receptor (25, 26), and Mpl (27). While multimeric forms of MPN mutant CRT have been described and implicated in Mpl activation and cytokine-independent cell growth (18, 28), the nature of the productive multimeric forms of CRT mutants that may trigger Mpl dimerization and activation has not been established. Here, based on the occurrence of novel cysteine residues in the CRT mutant C-termini (2, 3), we investigated the relevance of disulfide bond-mediated interactions in CRT multimerization in primary patient platelets and human cell lines expressing recombinant mutant CRT. In addition, we tested the relevance of non-covalent interactions relevant to CRT multimerization.

## Results

### Mutant CRTs form disulfide-stabilized multimers in MPN patient platelets and are detectable in MPN patient serum

To assess mutant CRT multimerization in primary cells, which has not previously been undertaken, platelets were purified from the blood of healthy donors or MPN patients with the known clinical characteristics summarized in Table S1 and analyzed by immunoblots. An antibody (anti-CRT(C_mut_)) was raised against the C-terminal 22 residues of mutant CRT, within the novel mutant-specific C-termini, to examine MPN mutant CRT expression and oligomerization. The specificity of the antibody was established using purified proteins (Figure S1). Anti-CRT(C_mut_) specifically detected proteins in platelet lysates from patients with CRT mutants but not from patients with JAK2 mutants or healthy control individuals (Figure 1A and Figure S2A, top panel). As expected, based on size, mutant CRT in samples 8744 and 4995-2 which correspond to CRT_Del4_ and CRT_Ins5_ mutations, migrate more slowly than the CRT_Del52_ samples in the same blots (Figure 1A and S2A, top panel). The levels of mutant CRT expression were somewhat variable between patients. For example, platelet lysates from consecutive blood collections from the patient 2648 consistently indicated high expression of the mutant CRT relative to other patients (Figure S2A for 2648-2).

**Fig. 1:**
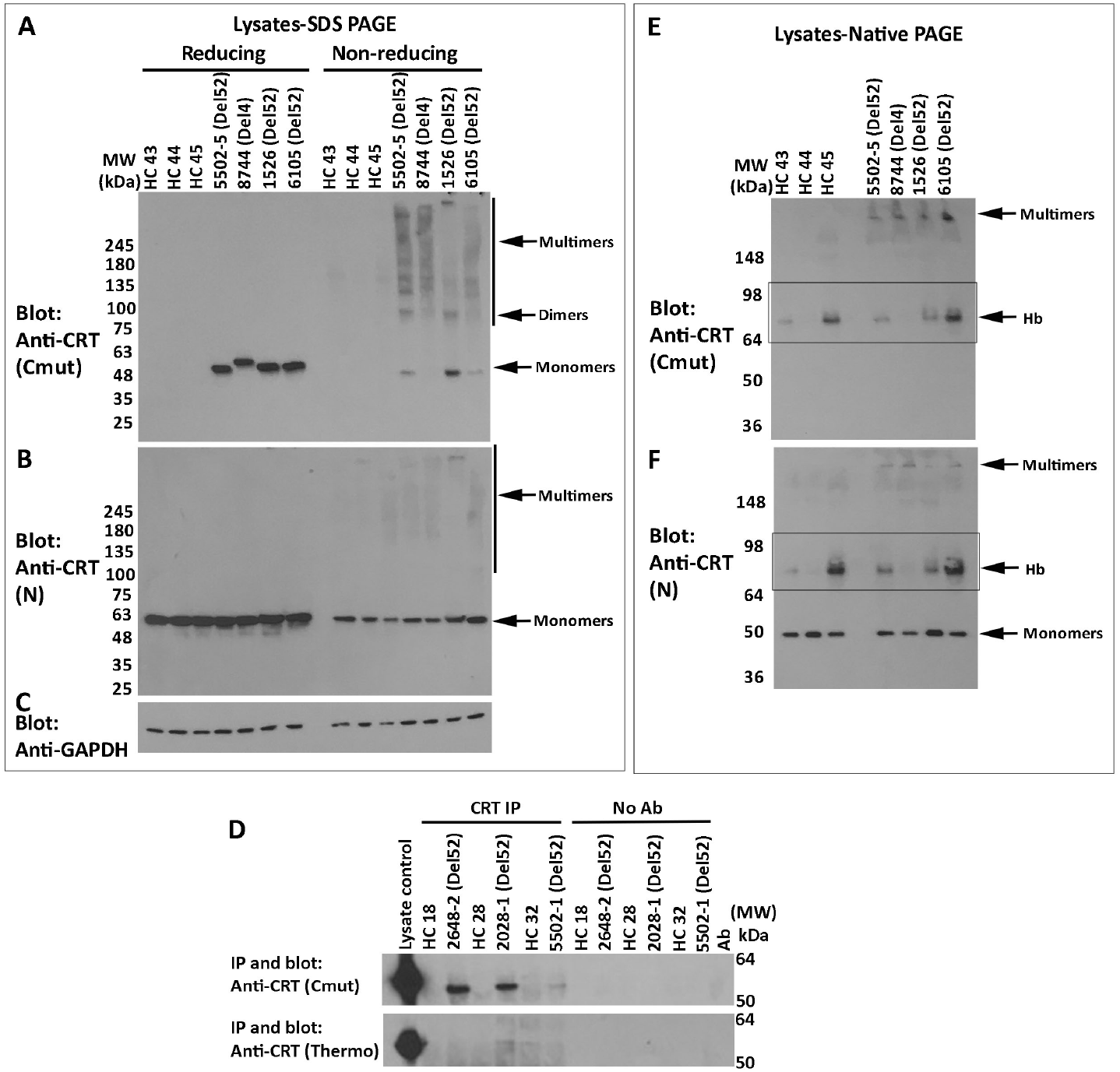
Mutant CRTs form disulfide-stabilized multimers in MPN patient platelets and are detectable in MPN patient serum. A-C) Lysates from MPN patients or healthy donor platelets were probed by SDS-PAGE under reducing or non-reducing conditions (using 4-20% gradient gels) followed by immunoblotting with anti-CRT(C_mut_) (A), anti-CRT(N) antibodies (B) or anti-GAPDH as control (C). The same lysates were loaded in two blots (A and B) and B was re-probed with anti-GAPDH (C). D) CRT secretion in healthy donor or MPN patient serum was examined by immunoprecipitation/immunoblotting analyses with anti-CRT (C_mut_) (top panel) or anti-CRT (Thermo Fisher) antibodies (bottom panel). No Ab lanes are controls in the absence of anti-CRT antibodies to assess non-specific precipitation. E and F) Representative immunoblots following native-PAGE (8% gels) of lysates from MPN patient platelets or healthy donor platelets, probed with the anti-CRT(C_mut_) (E) and anti-CRT(N) antibodies (F). The band indicated as multimers are over-represented in CRT mutant platelet lysates. Boxes indicate hemoglobin (Hb) contamination. In all panels, HC indicates healthy control samples; CRT mutant patient samples are indicated as Del52 or Del4 (characterized clinically as a 4 bp deletion in exon 9 resulting in a frameshift at K374, based on next generation sequencing. The predicted size of Del4 is 427 amino acids compared with 411 for Del52). See also Figure S1 for more information on the specificity of anti-CRT (C_mut_) and Figure S2 for additional replicate blots of platelet lysates.

In contrast to anti-CRT(C_mut_), anti-CRT(N), a commercial antibody from Cell Signaling Technology (CST) directed against the N-terminus of CRT, detected CRT in all samples (Figure 1B and Figure S2A, middle panels). As discussed below, in transfected HEK293T cells, anti-CRT(N) can distinguish CRT_Del52_ from CRT_WT_ based on the smaller size of CRT_Del52_. However, two distinct CRT bands were not readily detectable with anti-CRT(N) in lysates from MPN patient platelets (Figure 1B and Figure S2A and S2B, middle panels, except possibly for the most highly expressed samples such as 2648-2), indicating that the detected protein corresponds to CRT_WT_. The C-terminal frameshift in all MPN mutant CRT results in a loss of the ER retention KDEL motif, causing their secretion from cells (29–32) and indeed the mutants but not wild type CRT are detectable in patient serum by co-immunoprecipitation analyses (Figure 1D). Enhanced secretion is expected to render all MPN CRT mutants more difficult to detect than the wild type CRT using generic anti-CRT antibodies in platelet lysates.

Analyses of the platelet lysates from different patients on non-reducing gels indicated that the mutant CRT species are DTT-sensitive which was also most readily apparent in immunoblots with anti-CRT(C_mut_) for most samples, and with both anti-CRT(C_mut_) and anti-CRT(N) for the high-expressing samples 2648-3 and 8251-2 (Figures 1A-B and Figure S2B; reducing gels (+DTT) compared with non-reducing gels (-DTT)). Notably, a band consistent with a dimer of CRT (~100 KDa) was visualized in the non-reducing gels with anti-CRT(C_mut_) in several mutant CRT samples, in addition to several additional high molecular weight species (Figure 1A and S2B, top panel).

We also conducted immunoblots of the platelet lysates following native-PAGE. High molecular weight complexes, migrating close to the stacking gel, were clearly detectable with both anti-CRT(C_mut_) and anti-CRT(N) in the CRT mutant samples (Figure 1E-F and Figure S2C-D). Some platelet preparations show a band above the 64 kDa marker that likely represents a contamination of hemoglobin (Hb), based on the appearance of the same background in red blood cell (RBC) preparations (Figure 1E-F boxed; Figure S2D shows the RBC lysate), possible arising from RBC contamination in the platelet preparations. Thus, CRT mutants form disulfide-bonded high molecular weight complexes in patient platelets.

### The C-domain of CRT_Del52_ confers specificity for Mpl and forms disulfide-linked dimers

The novel C-domains of type-I and type-II MPN mutant CRT contain two or three cysteine residues (Figure 2A). To further study the C-domain multimerization and function, we generated C-domain fusions with the B1 domain of protein G (GB1; which is used to increase the yield and solubility of small proteins (33, 34)) and 6x histidine (his) tags. The C-domains of CRT_WT_, CRT_Del52_, and different truncation mutants of CRT_Del52_ (Figure 2A) were expressed as His-GB1-tagged proteins in HEK293T cells (Figure 2B). All constructs encoded the signal sequences of Cox2 to allow insertion into the ER lumen. Since the CRT_Del52_ mutation was the most frequent in our patient group (Table S1), our studies were largely focused on this mutant. Using the anti-His antibody, we found that the C-domain of CRT_Del52_ was expressed at higher levels than the C-domain of CRT_WT_ (Figure 2B, lane 1 compared to lane 2). Furthermore, truncations of CRT_Del52_ C-domain progressively reduced expression levels, with expression essentially undetectable for the C-domain of CRT_Del52Δ36_ (Figure 2B, lanes 2-6).

**Fig. 2:**
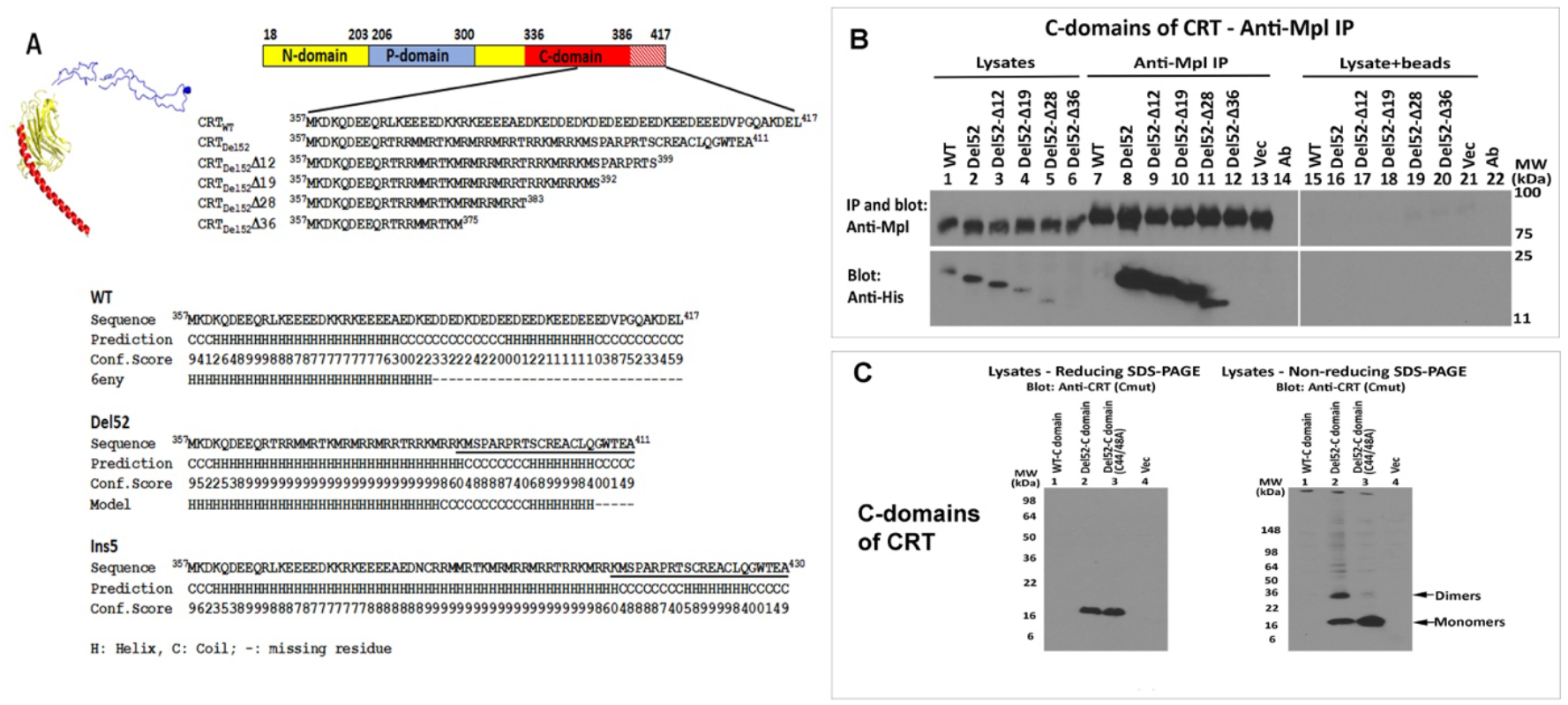
The C-domain of CRT_Del52_ confers specificity for Mpl and forms disulfide-linked dimers. A) Structure of CRT_WT_ (PDB ID: 6eny, subunit G (11)) with a globular lectin-like domain (yellow, residues 18-203 from N-domain and 301-335 from C-domain), a P domain (blue, residues 206-300), and an α-helical (residues 336-386) C-domain (red). The C-terminus of the C-domain (red stripes, residues 387-417) is not resolved in the structure. Sequences of CRT_Del52_ C-domain truncation constructs used in this study are shown. For C-domain expression, MPN-mutated sequences are extended by 10 residues at the N-terminus, to allow a common 10-mer sequence for the wild type and mutant C-domain constructs. The lower panel shows indicated C-domain sequences (line 1), secondary structure predictions performed using I-TASSER (41) (lines 2-3), the secondary structure based on PDB: 6eny or the CRT_Del52_ model discussed in this study (line 4). The sequence of the mutant-specific C-tail used to produce the anti-CRT(C_mut_) antibody is underlined. B) HEK293T cells were transiently transfected with plasmids encoding N-terminal His-GB1-tagged C-domains of CRT_WT_, CRT_Del52_, and truncated CRT_Del52_ C-domain constructs (sequences indicated in A), or a plasmid lacking CRT C-domain (Vec) along with N-terminal His-FLAG-tagged Mpl. Lysates from indicated cells were immunoprecipitated (indicated as IP) with anti-Mpl antibody or in the absence of anti-Mpl (marked as lysate+beads) and subsequent immunoblotting analyses were undertaken with the indicated antibodies. Data are representative of 3 independent experiments. C) HEK293T cells transiently transfected as indicated with plasmids encoding N-terminal His-GB1 tagged C-domains of CRT_WT_ (CRT_WT-C_), CRT_Del52_ (CRT_Del52-C_), cysteines mutated C-domain construct CRT_Del52-_ C(C44A/C48A) or a plasmid lacking CRT C-domain (Vec). Cell lysates from indicated cells were separated by SDS-PAGE under reducing (12.5% gels) (left panels) or non-reducing (4-20% gradient gels) (right panels) conditions and immunoblotted with indicated antibodies. Data are representative of 4 independent experiments. See also Figure S3 for additional data on Mpl specificity for full-length mutant CRT and Figure S4 for data on the inability of CRT_Del52_ C-domain constructs to induce Mpl proliferation.

We performed co-IP assays in HEK293T cells co-transfected with plasmids encoding Mpl and the various C-domain constructs. Binding interactions were observed between Mpl and the isolated C-domains of all the CRT_Del52_ constructs, except the poorly expressed CRT_Del52Δ36_, whereas binding between Mpl and the C-domain of CRT_WT_ was not observed (Figure 2B, lanes 7-12). The absence of C-domain bands in the lysate+beads controls suggests that binding of C-domains to Mpl is specific, as protein G contained on the beads is not bound by the C-domain constructs (Figure 2B, lanes 16-19).

The findings of Figure 2B parallel the previous findings of preferential interactions between the full-length versions of MPN mutant CRT and Mpl in the steady state (12, 13, 18-22). Correspondingly, in an anti-Mpl co-IP with HEK293T cells transfected with plasmids encoding Mpl and full-length mutant CRT constructs, binding interactions were readily detectable between CRT mutants and Mpl using anti-CRT(C_mut_) (Figure S3A, top and middle panels for Mpl and anti-CRT(C_mut_) blots). On the other hand, CRT_WT_ signals (probed with anti-CRT(Thermo), a commercial polyclonal antibody directed against the whole protein as immunogen; Thermo Fisher), were essentially undetectable in the steady state, following anti-Mpl co-IPs (Figure S3A, lower panel). Wild type but not mutant protein over-expression is detectable with anti-CRT(Thermo) (Figure S3A, compare lanes 1-3 with lane 4). Again, as noted above, anti-CRT(C_mut_) allows for more sensitive detection of the mutants, and additionally, it is possible that anti-CRT(Thermo) epitopes reside within the C-terminus. Preferential binding between mutant CRT and Mpl was also detectable in patient platelets (Figures S3B and S3C).

In lysates of transfected HEK293T cells, a band consistent with the size of a C-domain dimer was detected under non-reducing conditions, for the CRT_Del52-C_ but not its cysteine mutant, CRT_Del52-C(C44A/C48A)_ (Figure 2C, right panel, lanes 2 and 3). Thus, the mutant C-domain, when expressed on its own, is capable of forming disulfide-linked dimers.

For further functional assessments of the C-domain constructs, we first used retroviral infections to stably express Mpl in Ba/F3 cells, and further expressed CRT_Del52_ or its C-domain or cysteine mutated C-domain. Binding interactions between Mpl and the CRT_Del52_ full-length and C-domain constructs were observed in Ba/F3 Mpl cells (Figure S4A), as was observed in HEK293T cells (Figure 2B). The efficiency of Mpl binding to the C-domain constructs appears to be at least as high as that for full-length CRT_Del52_ (Figure S4A; lane 2 compared with lane 4), even though expression of the C-domain constructs is not directly detectable in Ba/F3 lysates. MPN mutant CRT constructs are known to induce proliferation of Ba/F3 cells in a Mpl-dependent and cytokine-independent manner (13, 22), as also shown in Figure S4B. Consistent with the previous findings (19), the CRT_Del52_ C-domain alone is insufficient to mediate proliferation (Figure S4B). Thus, while the CRT_Del52_ C-domains confer Mpl binding specificity and dimerize (Figure 2), additional interactions mediated by CRT_Del52_ are required for functional interactions with Mpl (Figures S4). Notably, the CRT_Del52_ C-domain has higher stability than the wild type C-domain in HEK293T cells, based on relative expression levels in lysates, both in the presence and in the absence of Mpl (Figure 2B and S4C respectively), consistent with the higher predicted helical content of the CRT_Del52_ C-domain (Figure 2A).

### Truncations of C-terminal cysteines of full-length CRT_Del52_ alter but do not abrogate disulfide-linked interactions

To ask if C-domain cysteines were sufficient for full-length CRT_Del52_ multimerization, we generated the successive truncations of the C-terminal sequences (Figure 2A) within full-length CRT_Del52_ and expressed those constructs as N-terminal His and GB1-tagged (his-GB1) proteins. CRT_Del52Δ12_ and CRT_Del52Δ19_ were expressed at low levels and their protein loads had to be increased to achieve similar protein expression as the other constructs (Figure 3A; note varying intensities of the endogenous CRT bands). Furthermore, in non-reducing SDS gels (Figure 3B), only CRT_WT_ and CRT_Del52Δ36_ were largely monomeric. Bands consistent with the size of dimers/multimers were observed for CRT_Del52_, CRT_Del52Δ12_, CRT_Del52Δ19_, and CRT_Del52Δ28_, even though all of the truncation constructs of CRT_Del52_ lacked the two C-terminal cysteines. However, the specific band indicated as dimers for CRT_Del52_ was absent in CRT_Del52Δ12_, CRT_Del52Δ19_, and CRT_Del52Δ28_, and instead slower mobility bands were observed (Figure 3B, lane 3, compared to lanes 4-6). These findings suggested that the presence of two C-terminal cysteines in CRT_Del52_ contribute to the induction of distinct disulfide-linked species. Additionally, the proportion of monomer bands to total CRT bands progressively increases with increased truncation size (quantification of 3B is shown in Figure 3C). In native gels, only multimers were detectable for CRT_Del52Δ12_ and CRT_Del52Δ19_, monomers and multimers were detected for CRT_Del52Δ28_, whereas CRT_Del52Δ36_ migrated largely as monomers (Figure 3D). Together the findings of Figure 3 suggested that novel C-domain cysteines and additional cysteines contribute to the formation of CRT_Del52_ disulfide-linked dimers. Further, the CRT_Del52Δ28_ truncation is needed to partially destabilize CRT_Del52_ multimers and the CRT_Del52Δ36_ truncation is needed to fully destabilize CRT_Del52_ multimers, suggesting that both covalent and non-covalent interactions mediated by the C-domains contribute to multimer formation.

**Fig. 3:**
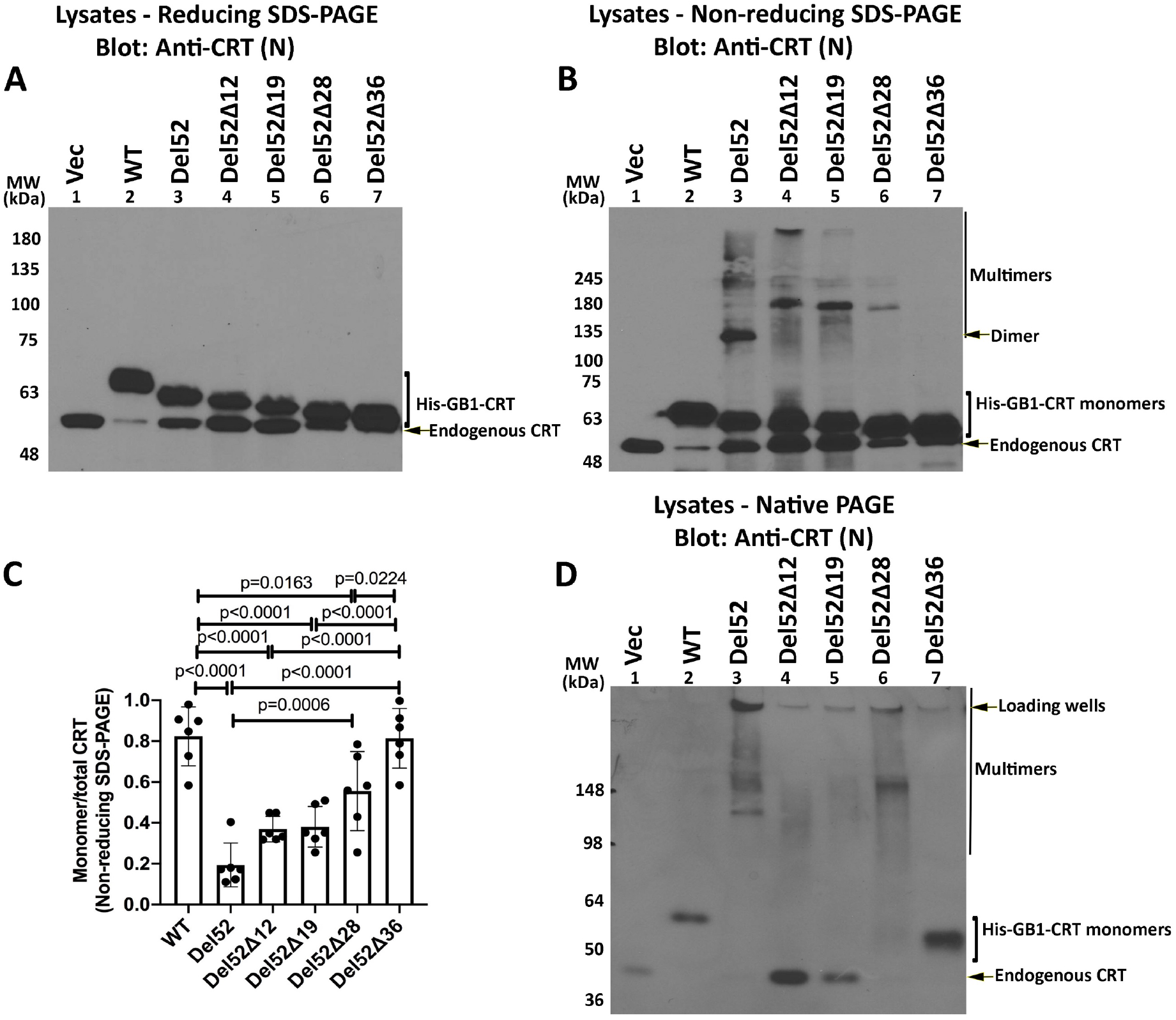
Truncations of C-terminal cysteines of full-length CRT_Del52_ alter but do not abrogate disulfide-linked interactions. HEK293T cells were transiently transfected with plasmids encoding N-terminal His-GB1 tagged full-length or C-terminally truncated CRT_Del52_ constructs. Cell lysates from transfected cells were separated by SDS-PAGE under reducing (10% gels) (A) or non-reducing (10% gels) (B) conditions or by native-PAGE (4-20% gradient gels) (D) and immunoblotted with the anti-CRT(N) antibody. Different amounts of lysates were loaded to achieve similar protein expression of different truncated constructs (CRT_WT_ (0.5 μg lysates), CRT_Del52_ (5 μg lysates), CRT_Del52Δ12_ (18 μg lysates), CRT_Del52Δ19_ (18 μg lysates), CRT_Del52Δ28_ (3 μg lysates), CRT_Del52Δ36_ (1.8 μg lysates), or a plasmid lacking CRT (Vec) (10 μg lysates). The endogenous CRT band serves as the lysate loading controls. Species consistent with the size of endogenous CRT, His-GB1-CRT monomers, dimers, multimers and loading wells are indicated. Quantification of CRT monomer/monomer+multimer (total) bands from B is shown in C, averaged over 6 independent blots from 5 independent transfections. Data show mean ± SD, with statistical significance assessed via ordinary-one-way ANOVA.

### Disulfide-linked CRT_Del52_ dimer and multimer formation is C-domain and N-domain dependent

To further elucidate the mode of CRT_Del52_ multimerization, we generated various cysteine mutants of CRT_Del52_ as untagged constructs and examined their multimerization in transfected HEK293T cells. Anti-CRT(C_mut_) did not detect wild type CRT expressed in HEK293T cells and was specific for the mutants (Figure 4A). As noted above, CRT_Del52_ formed dimers and higher order species, which were detected in immunoblots under non-reducing conditions (Figure 4B, lanes 3-4). CRT_Del52-2CA_ (CRT_Del52(C400A/C404A)_, that lacks the novel cysteines in the mutant C-terminus) formed fewer higher order multimer structures and more lower order structures compared with CRT_Del52_ (compare Figure 4B lanes 5 and 6, which had similar protein loads as shown in Figure 4A, lanes 5 and 6, and quantification in 4C). Notably, the bands indicated as dimers (which migrated at approximately 100 kDa, the expected size for Del52 dimers) were more intense but migrated more slowly for CRT_Del52-2CA_ compared to the corresponding CRT_Del52_ construct (Figure 4B, lanes 3-4 compared with 5-6), suggesting that the CRT_Del52_ dimers are rendered more compact by the presence of the two C-terminal cysteines, C400 and C404. CRT has a single disulfide bond between C105 and C137 within its globular domain. C163, the only free cysteine in the wild type CRT that is also present in MPN-linked CRT mutants. Ala-substitution of this residue in the CRT_Del52-CA_ (CRT_Del52(C163A)_) mutant resulted in a dimer and multimer pattern similar to CRT_Del52._ Notably, the intensities of bands indicated as monomers and dimers were stronger for CRT_Del52-CA_ compared with CRT_Del52_, whereas bands indicated as multimers were stronger for CRT_Del52_ compared with CRT_Del52-CA_ (Figure 4B, lanes 9-10 compared with lanes 3-4 and quantification in Figure 4C), under conditions of similar protein loads (Figure 4A, lanes 9-10 compared with lanes 3-4). Additionally, monomer species were predominant for the triple cysteine mutant CRT_Del52-_ 3CA (CRT_Del52(C163A/C400A/C404A)_) indicating that all three mutations were needed to significantly inhibit the formation of disulfide-linked dimers and multimers (Figure 4B, lanes 7-8 and Figure 4C). C105 and C137 are the only remaining cysteines in CRT_Del52-3CA_, residues that form a disulfide bond in CRT structures (6, 7). The disulfide linked dimers and multimers still observable with CRT_Del52-3CA_ likely correspond to low efficiency thiol-disulfide exchange between the monomer subunits of dimers/multimer.

**Figure 4:**
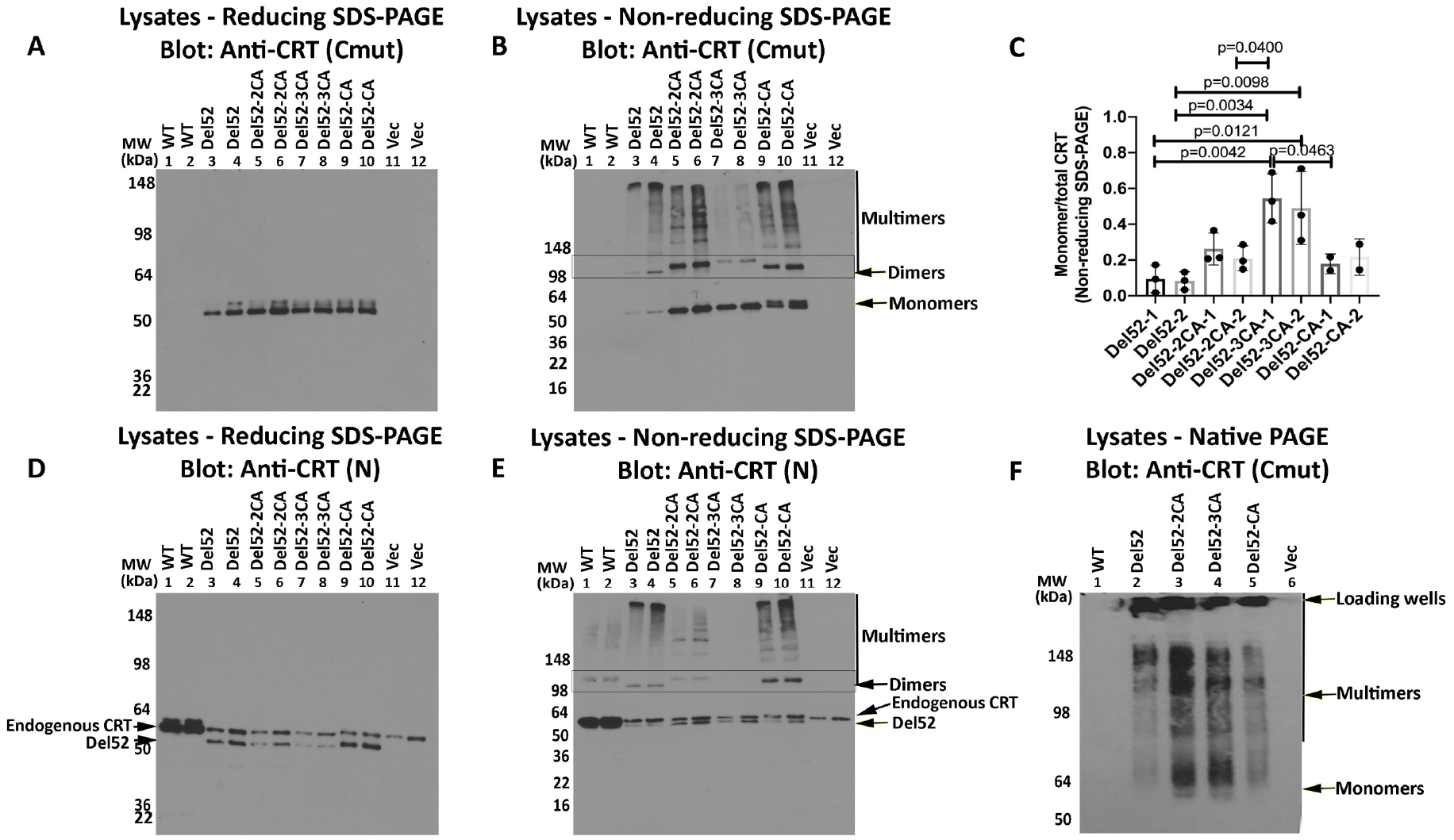
Disulfide-linked CRT_Del52_ dimer and multimer formation is C-domain and N-domain dependent. HEK293T cells were transiently transfected with plasmids encoding untagged full-length CRT_WT_, CRT_Del52_, CRT_Del52-2CA_ (CRT_Del52(C400A/C404A)_), CRT_Del52-3CA_ (CRT_Del52(C163A/C400A/C404A)_), CRT_Del52-CA_ (CRT_Del52(C163A)_) or plasmid lacking CRT (Vec). Cell lysates from indicated cells with two lysate amounts (A-B, D-E) were separated by SDS-PAGE under reducing (8% gels) (A, D) or non-reducing (4-20% gradient gels) (B, E) conditions and immunoblotted with indicated antibodies. Full gel panels are shown. Data are representative of three independent experiments for CRT_WT_, CRT_Del52_, CRT_Del52-2CA_, CRT_Del52-3CA_ and two independent experiments for CRT_Del52-CA_. Species consistent with the size of CRT monomers, dimers and multimers are indicated. In panel D where CRT_WT_ and CRT_Del52_ constructs are resolved, the migration position of each are indicated as endogenous CRT and CRT_Del52_. C) Quantifications of blots from B to calculate monomer/monomer+multimer (total) mutant CRT following non-reducing SDS-PAGE are shown in C, averaged over 3 independent experiments (two protein loads each). Consistent ratios are quantified at two different protein loads of each construct (labeled as 1 or 2). Data show mean ± SD, with statistical significance assessed via ordinary-one-way ANOVA. F) Cell lysates from indicated transfected cells were separated by native-PAGE (8% gels) and immunoblotted with indicated antibody. Bands corresponding to CRT_Del52_ monomers, multimers and loading wells are indicated. See also Figure S5 for disulfide-linked multimer formation by CRT_Ins5._

Parallel immunoblots under reducing conditions with anti-CRT(N) indicated that all CRT_Del52_ constructs migrate more rapidly than CRT_WT_, consistent with their smaller size (Figure 4D). Additionally, bands indicated as dimeric and high order oligomeric CRT structures were detectable under non-reducing conditions similarly to those detected with anti-CRT(C_mut_) (Figure 4E). Slower migration of CRT_Del52-2CA_ dimer bands than CRT_Del52_ (Figure 4E, lanes 3-4 compared with 5-6), and higher representation of dimer bands relative to higher order multimers for CRT_Del52-CA_ compared with CRT_Del52_ were again rather noticeable (Figure 4E, lanes 3-4 compared with 9-10). Finally, disulfide-linked multimeric structures were poorly detected for the triple mutant CRT_Del52-3CA_. Together, these finding implicate C-terminal cysteines (C400 and C404) and C163 in disulfide-mediated dimerization and multimerization of CRT_Del52_ mutant.

In native-PAGE gels, however, all four CRT_Del52_ constructs formed higher order species (Figures 4F). Nonetheless, signals corresponding to monomeric species were readily detectable only in the CRT_Del52-2CA_ and CRT_Del52-3CA_ lysates (Figure 4F). We concluded from these analyses that both covalent and non-covalent interactions contribute to CRT_Del52_ multimerization. Together with the truncation mutant data (Figure 3), these findings indicate that disulfide-dependent interactions contribute to dimer and multimer stability, but that loss of S-S bonds is not sufficient to fully block multimer formation.

Reducing and non-reducing SDS-PAGE analyses indicated the formation of disulfide-linked oligomers for CRT_Ins5_ constructs, similarly to that observed for CRT_Del52_, although the overall pattern was more complex than that of CRT_Del52_ because of the presence of an additional C-terminal cysteine in CRT_Ins5_ mutant (Figure 2A and Figure S5). Further mutational studies were focused on CRT_Del52_ mutant.

### A working model for a CRT_Del52_ dimer including ionic interactions mediated by the N-domain

We generated a model for monomeric CRT_Del52_ based on the high resolution crystal structure of the human CRT (35) for the N-domain and the proximal part of C-domain (residues 19-203 and 303-366, excluding the P-domain). The C-domain α-helix was further extended to K386 using the structure of the human major histocompatibility complex class I peptide-loading complex which includes CRT (11). The distal C-terminal segment (residues 399-406) of CRT_Del52_ containing C400 and C404 was modeled as a 2-turn α-helix based on secondary structure predictions by I-TASSER (Figure 2A) while the connecting loop (residues 387-398) was modeled in an extended conformation.

We next examined various crystal structures of CRT for dimerization modes that could account for the experimental data (Figures 1–4). Among the structures (see supplemental methods for details), the crystal structure of a 10-mer of human CRT (D71K mutant; PDB ID: 5lk5) (35) revealed stable dimers formed through tight packing of antiparallel α-helices from C-domains (Figure S6A, “C-C” dimer) or through intermolecular ionic interactions between N-domain loop residues 160-167 (Figure S6B, “N-N” dimer). This N-domain loop contains C163, which, based on our data (Figure 4), is involved in disulfide-mediated dimerization. In the crystal structure of the “N-N” dimer, two C163 are not in direct contact, but could move closer to each other and form a disulfide bond following minor loop rearrangements. In addition, the C-termini of molecules in the “N-N”-dimer are much closer to each other than in the “C-C”-dimer, and, therefore, the “N-N”-dimer can be additionally connected through two intermolecular disulfides between C400 and C404 of CRT_Del52_ (Figures 2–4). Two negatively charged D165 residues from both subunits of the “N-N” dimer form hydrogen bonds and four intermolecular ionic pairs with positively charged K142 and R162 from the opposite subunits. Closer examination of the “N-N” dimerization mode of CRT_Del52_ mutants (Figure 5A) allows prediction of two additional symmetrical “N-C” dimerization interfaces that could be formed between N-domain of one molecule and C-domain of another molecule. A slight decrease of the α-helix kink at A352 would move the C-terminal part of the α-helix (residues 366-383) closer to N-domain glycan recognition site. As a result, dimer-stabilizing ionic and hydrophobic interactions may be formed between positively charged and non-polar residues from C-domain of one molecule and negatively charged and aromatic residues from the N-domain glycan-binding site of the second molecule. The existence of ionic or hydrophobic interactions between these residue pairs may explain the significant role of residues 376-383 for multimerization of CRT_Del52_ constructs, as their truncation in CRT_Del52Δ36_ eliminated oligomer formation (Figure 3). Thus, the “N-N” dimerization mode is compatible with our experimental data (Figures 2–4). Furthermore, CRT_Del52_ tetramers and larger oligomers can be easily formed by combining both types of dimerization modes: through C-domain helix-helix interface and N-domain loop-loop interface (Figure S6C).

**Fig. 5:**
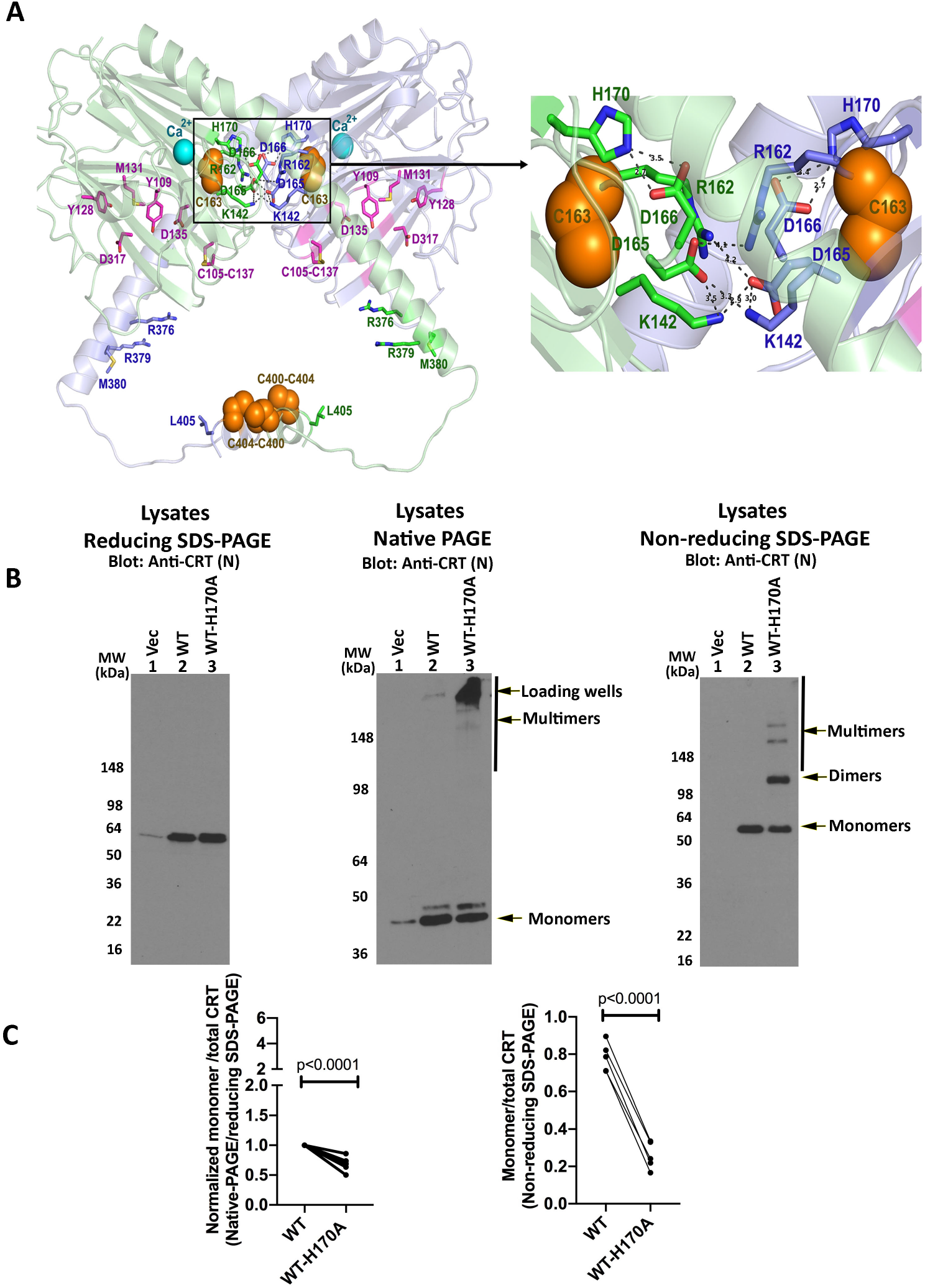
A working model for a CRT_Del52_ dimer and assessment of H170-regulated wild type CRT multimerization. A) A molecular model of CRT_Del52_ dimer is primarily based on the crystal structure of the 10-mer complex of CRT D71K mutant (PDB ID: 5lk5, subunits E and G (35)). Subunit 1 is colored light green and subunit 2 light blue. Each subunit contains a globular N-domain and a C-domain comprising long proximal and short distal α-helices connected by a 14-residue loop. The P-domain is omitted. Ca^2+^-ions bound to the high-affinity site are shown by cyan spheres. Residues from the carbohydrate recognition site (7) are shown by purple sticks (for Cα-atoms). Cysteines participating in the formation of predicted intermolecular disulfide bonds (C163-C163 and two C400-C404) are shown by orange spheres. To form a C163-C163 disulfide bond, loops have to be rearranged to decrease the distance between cysteines. Dimers are also stabilized by intermolecular hydrogen bonds (shown by gray dashes, distances < 3.0 Å) and ionic interactions (shown by gray dashes, distances < 4.5 Å) that are formed between charged residues from the N-domain dimerization interface: D165, K142, and R162 (residues are shown by sticks colored green for subunit E and blue for subunit G). Inset highlights interactions at the contact interface. Molecular graphic representations were generated by PyMOL. See also Figure S6 for models of the structural organization of CRT dimers into higher order multimers. B) HEK293T cells were transiently transfected with plasmids encoding untagged full-length CRT_WT_, CRT_WT-H170A_ or plasmid lacking CRT (Vec). Cell lysates from indicated cells were separated under reducing SDS-PAGE (8% gel, left panel) or native-PAGE (8% gel, middle panel) or non-reducing SDS-PAGE (4-20% gradient gel, right panel) conditions and immunoblotted with anti-CRT (N) antibody. C) Left panel: Monomer bands for each construct from the native-PAGE immunoblots were quantified and normalized relative to the total CRT signal from parallel SDS-PAGE immunoblots, averaged over 6 independent blots from 4 independent transfections. Data show mean ± SD, the normalized signals were log transformed and the statistical significance was assessed via one sample t-test. Right panel: Quantifications of monomer/monomer+multimer (total) mutant CRT bands from non-reducing SDS-PAGE immunoblots, averaged over 5 independent blots from 4 independent transfections. Data show mean ± SD and the statistical significance was assessed via a paired t-test.

We previously showed that the H170A mutant of murine CRT (H153A in mature protein numbering) forms dimers at enhanced levels, visualized by native-PAGE of the purified protein (36). H170 is at predicted dimer interface in the model shown in Figure 5A and forms intramolecular interactions with D166. We found that H170A mutations of wild type human CRT also induce disulfide-linked dimers and multimers, visualized in transfected HEK cells (Figures 5B and 5C), indicating the potential relevance of the dimer model of Figure 5A to multimeric interactions in both wild type CRT and its MPN mutants. We and others have also previously reported that disulfide-linked dimers of CRT are induced by heat-shock, calcium depletion and other destabilizing conditions (37, 38).

Based on the dimer model of Figure 5A, in addition to the covalent interactions, CRT_Del52_ dimers are stabilized by two sets of D165-K142 and D165-R162 salt bridges at the globular domain dimer interface. The D165K mutations alone (CRT_Del52-D165K_) did not significantly influence the oligomerization potential of CRT_Del52_ (Figure S7). However, the combination of D165K and CRT_Del52-3CA_ mutations induced more monomeric species in the CRT_Del52-3CA/D165K_ mutant compared with CRT_Del52-3CA_ in native blots, but not in non-reducing blots (Figures 6A and 6B), indicating the impact of combined disruptions of both dimer interfaces. This was not observed with the D166K mutation, for which enhanced levels of disulfide-linked species were induced (with both CRT_Del52-3CA/D166K_ and the double mutant CRT_Del52-3CA/D165KD166K_ (CRT_Del52-3CA-2DK_) compared with CRT_Del52-3CA_ (Figures 6A and 6B). Since the only remaining cysteines in all CRT_Del52-3CA_ constructs are C105 and C137, which form an intramolecular disulfide in wild type CRT (6, 7), these findings suggest that repulsion between several interacting positively charged residues in the D166K mutant following disruption of its interaction with H170, induce rearrangements that bring the C105-C137 disulfide from one subunit of the N-N dimer closer (from 18 Å to 5 Å) to that from the other subunit to enhance intermolecular disulfides via thiol-disulfide exchange. Correspondingly, the H170A mutation on the CRT_Del52-3CA_ background also induces disulfide-linked dimers and multimers (Figures 6C and 6D), that are expected to correspond to enhanced intermolecular disulfides mediated by C105-C137 rearrangements. Overall, the findings of Figure 6 support the relevance of the dimer model of Figure 5A to Del52 multimerization, and suggest that MPN mutations exploit a natural dimerization interface of CRT known to be induced by ER stress conditions.

**Fig. 6:**
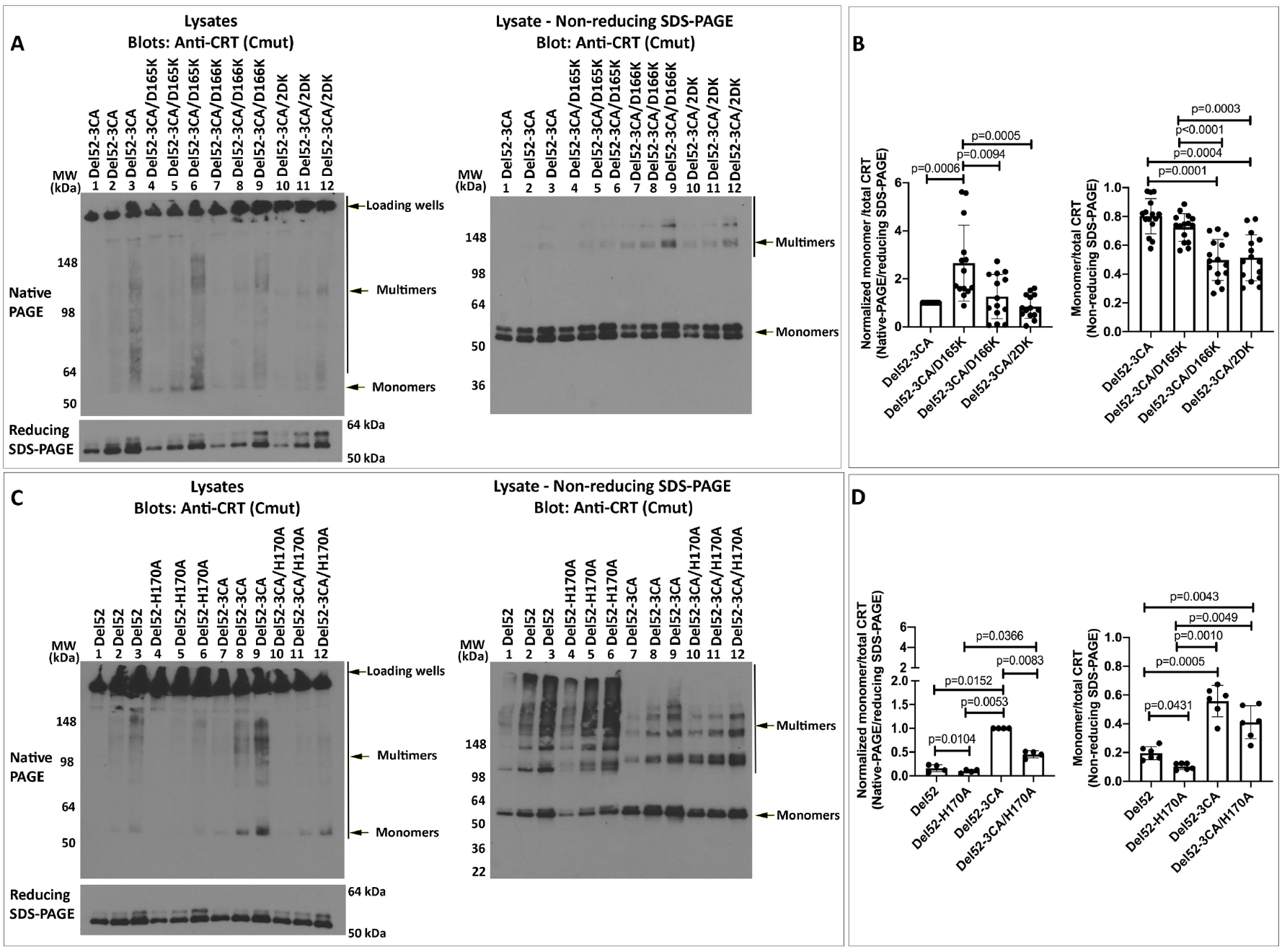
N-domain dimer interface residues influence CRT multimerization. HEK293T cells were transiently transfected as indicated with plasmids encoding untagged full-length CRT_Del52-3CA_ (CRT_Del52(C163A/C400A/C404A)_), CRT_Del52-3CA/D165K_ (CRT_Del52(C163A/C400A/C404A/D165K)_), CRT_Del52-3CA/D166K_ (CRT_Del52(C163A/C400A/C404A/D166K)_), and CRT_Del52-3CA/2DK_ (CRT_Del52(C163A/C400A/C404A/D165K/D166K)_), CRT_Del52-H170A_ or CRT_Del52-3CA/H170A_ (CRT_Del52(C163A/C400A/C404A/H170A)_). Cell lysates from indicated cells were separated by native-PAGE (8% gels) (A and C, top left panels) or SDS-PAGE under reducing (8% gels) (A and C, bottom left panel) or non-reducing (4-20% gradient gels) (A and C, right panels) and immunoblotted with the anti-CRT (C_mut_) antibody. B and D) Left panels: Mutant CRT monomer bands from native-PAGE immunoblots in panels A and C were normalized relative to the corresponding total CRT signal from the reducing SDS-PAGE immunoblots. Data show mean ± SD, the normalized signals were log transformed and the statistical significance was assessed via RM one-way ANOVA analysis. B and D) Right Panels: Quantification of CRT monomer/ monomer+multimer (total) bands from the non-reducing SDS-PAGE immunoblots. Data show mean ± SD and the statistical significance was assessed via RM one-way ANOVA analysis. Data were averaged over 5 independent blots from 5 independent transfections (B) or 2 independent blots from 2 independent transfections (D), each with 2-3 protein loads. See also Figure S7 for multimer formation by CRT_Del52-D165K_ and CRT_Del52-D166K_.

### Large C-domain truncations or combined N-domain and C-domain dimer interface mutations are required to abrogate CRT_Del52_-mediated cell proliferation

Ba/F3-Mpl cells were transduced with viruses encoding the series of untagged C-domain truncated constructs of CRT_Del52_ and its point mutants to compare their proliferation-inducing activities. We observed reduced abilities of CRT_Del52Δ12_ and CRT_Del52Δ19_ to mediate Ba/F3 cell proliferation (although statistically non-significant), while CRT_Del52Δ28_, CRT_Del52Δ36_ were unable to promote cell growth, similar to CRT_WT_ (Figure 7A; all constructs shown in Figure 7 are untagged). These results deviate from those of Elf et al (20), where the transforming capacity of CRT mutant was abolished only after the most severe truncation of its C-terminus in CRT_Del52Δ36_. We could not unambiguously detect expression of the truncated untagged CRT constructs over interfering background bands in two independent sets of retroviral infection of Ba/F3-Mpl cells (data not shown). Thus, we re-assessed expression and functional activities of N-terminal histidine and GB1-tagged versions in Ba/F3-Mpl cells following plasmid nucleofections (Figure S8). Proliferation mediated by tagged versions of CRT_Del52Δ28_ and CRT_Del52Δ36_ was again impaired, under conditions where expression of both those constructs was detectable at higher levels than of CRT_Del52_ (Figure S8). Thus, while removal of the novel C-terminal cysteines has a small effect on CRT_Del52_ mediated proliferation, a larger truncation is needed to completely abrogate CRT_Del52_ mediated proliferation (Figure 7A and S8).

**Fig. 7:**
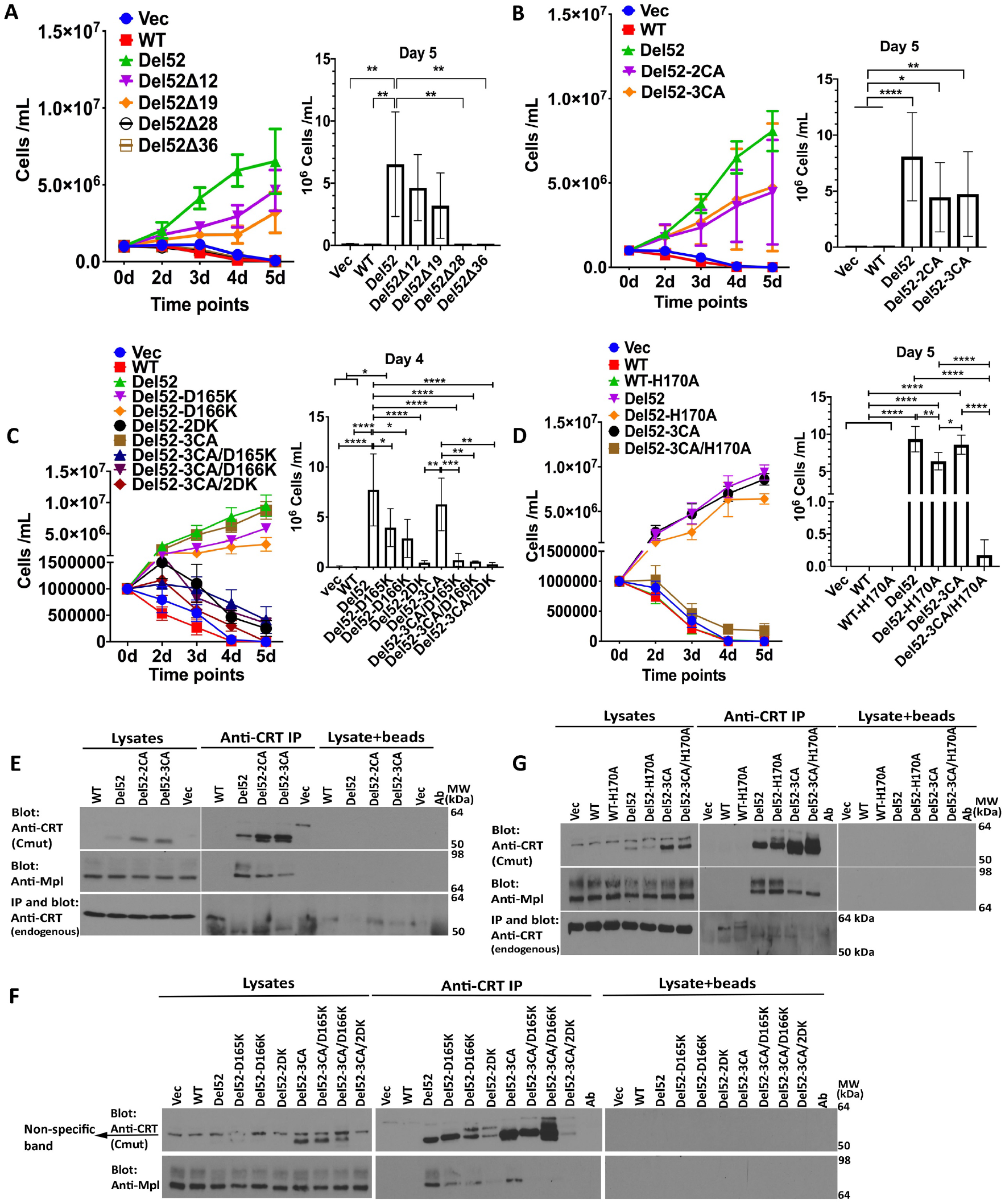
Large C-domain truncations or combined N-domain and C-domain dimer interface mutations are required to abrogate CRT_Del52-_mediated cell proliferation. Ba/F3-Mpl cells as specified were transduced with retroviruses (Ba/F3-Mpl) encoding full-length untagged CRT_WT_, CRT_Del52_, its truncation constructs (CRT_Del52Δ12_, CRT_Del52Δ19_, CRT_Del52Δ28_, CRT_Del52Δ36_), point mutants (CRT_Del52-2CA_ (CRT_Del52(C400A/C404A)_), CRT_Del52-3CA_ (CRT_Del52(C163A/C400A/C404A)_), CRT_Del52-D165K_, CRT_Del52-D166K_, CRT_Del52-2DK_ (CRT_Del52(D165/166K)_), CRT_Del52-3CA/D165K_ (CRT_Del52(C163A/C400A/C404A/D165K)_), CRT_Del52-3CA/D166K_ (CRT_Del52(C163A/C400A/C404A/D166K)_), CRT_Del52-3CA/2DK_ (CRT_Del52(C163A/C400A/C404A/D165K/D166K)_), CRT_WT-H170A_, CRT_Del52-H170A_, CRT_Del52-3CA/H170A_ (CRT_Del52(C163A/C400A/C404A/H170A)_)), or control viruses and used for the subsequent analyses. A-D) Cytokine-independent proliferation of Ba/F3-Mpl cells. Cell proliferation was measured as described in Figure S4 legend. Data are averaged from three separate viral transductions of Ba/F3-Mpl cells and a total of 5 independent proliferation experiments (A), 8 separate viral transductions of Ba/F3-Mpl cells, and a total of 10-13 independent experiments (B), 2-3 separate viral transductions of Ba/F3-Mpl cells, and a total of 3-6 experiments (C) or two separate retroviral transductions of Ba/F3-Mpl cells, and a total of 5 independent experiments (D). Mean ± SD is shown, with statistical significance assessed via ordinary-one-way ANOVA from the indicated days of proliferation assay. Statistically significant means are indicated as * *p* <0.05, ** *p* <0.01, *** *p* <0.001, and **** *p* <0.0001. E-G) Lysates from Ba/F3-Mpl cells expressing indicated constructs or control cells expressing Mpl alone (Vec) were directly loaded for immunoblotting analyses (labeled as lysates) or immunoprecipitated (indicated as IP) with anti-CRT(C_mut_) antibody (for CRT_Del52_ and its variants) or with anti-CRT(Thermo) antibody (for CRT_WT_) and subsequent immunoblotting was undertaken with the indicated antibodies. Results are representative of 3 (E and F) or 4 (G) independent experiments. Non-specific interactions in the absence of primary antibody are shown by the lysate+ beads lanes. See also Figure S8 for proliferation induction by His-GB1-tagged versions of the CRT_Del52_ truncation mutants.

Both CRT_Del52-2CA_ and CRT_Del52-3CA_ showed non-significant reductions in the ability to induce cytokine-independent proliferation of Ba/F3-Mpl cells compared with CRT_Del52_, (Figure 7B), despite the higher expression of the mutant (Figure 7E, top panel, lysate lanes). Notably, we also observed decreased binding of CRT_Del52-2CA_ and CRT_Del52-3CA_ to Mpl relative to CRT_Del52_ despite the higher expression of the mutants (Figure 7E, lanes marked as IP). There was also a non-significant reduction in Ba/F3-Mpl cell proliferation induced by CRT_Del52-D165K_ and CRT_Del52-D166K_ compared with CRT_Del52_ (Figure 7C). Notably, the combination of D165K or D166K or both with the CRT_Del52-3CA_ mutation resulted in marked abolishment of cell proliferation (Figure 7C). The impaired abilities of CRT_Del52-3CA/D165K_ and CRT_Del52-3CA/D166K_ to mediate cytokine-independent cell proliferation correlated with almost complete impairment in Mpl binding (Figure 7F, IP lanes). Parallel results were obtained with CRT_Del52-3CA/H170A_ (Figure 7D and 7G).

Among all mutants, the expression level of the combined mutant CRT_Del52-3CA/2DK_ (3CA+D165K+D166K) was rather low. Thus, the impaired cell proliferation induced by this mutant could be partially caused by its low expression. In contrast, the expression of CRT_Del52-3CA_, CRT_Del52-3CA/D165K_ and CRT_Del52-3CA/D166K_, CRT_Del52-3CA/H170A_ mutants were higher than of CRT_Del52_ (Figures 7F and 7G, top panels), indicating an enhanced stability of these mutants. Therefore, loss of stability could not explain their functional loss.

## Discussion

In this work, we observed that the presence of the novel C-terminal domain in MPN mutant CRT induces CRT dimerization and formation of higher order oligomers in pathologically relevant MPN patients-derived platelets (Figure 1), and in transfected HEK293T cells (Figures 2–4). Absence of oligomeric bands under disulfide-reducing conditions indicates that the intermolecular disulfides stabilize multimers both in transfected cells and in MPN patient-derived platelets.

The data presented here are consistent with the model of dimers of CRT_Del52_ with two dimerization interfaces, the first involving interactions between the distal parts of C-terminal tails and the second involving association of globular domains via loop residues 160-167 (Figure 5A). The novel C-domains of the MPN-linked CRT mutant, CRT_Del52_, contains two cysteine residues, C400 and C404, (Figure 2A), whose mutations to alanines abrogate the formation of dimers of isolated C-domains (Figure 2C). In addition, C163 residue from the globular N-domain contributes to the formation of intermolecular disulfides. Indeed, a combination of mutations of two C-terminal cysteines and the C163A mutation in the full-length CRT_Del52_ increases the fraction of monomers (Figures 4B, 4C and 4F). Based on available crystal structures of CRT oligomers, we propose a model of CRT_Del52_ dimer that is consistent with our experimental data, wherein dimer stabilizing interactions occur through N-domains (C163-C163 disulfide and four ionic bridges involving D165) and through C-tails cross-linked by two C400-C404 disulfides (Figure 5A, and related discussion). The combinations of D165K with the triple cysteine mutant in CRT_Del52-3CA_ significantly increases the fraction of the monomeric form of the protein. This is not observed with CRT_Del52-3CA/D166K_, which appears to induce structurally modified dimers (Figure 6B), which nonetheless are not compatible with Mpl binding and activation. Combined mutations that interfere with both dimerization interfaces (CRT_Del52-3CA/D165K_, CRT_Del52-3CA/D166K_ and CRT_Del52-3CA/H170A_) completely abrogate CRT_Del52-_induced cell proliferation (Figures 7C and 7D).

Previous findings have indicated preferential binding of full-length CRT mutants to Mpl in transfected cells (12, 13, 18-22). Here, we observed the binding of Mpl not only to the full-length CRT_Del52_ (Figure S3A) but also to its isolated C-terminal domain (Figures 2B and S4). This is direct evidence that the novel C-terminus of CRT_Del52_ confers the Mpl binding specificity. Cysteine residues within the mutant C-domains are not absolutely required for Mpl binding either to the full-length CRT_Del52_ or to its C-domain (Figures 2B, 7E, and S4). However, mutations C400A/C404A in CRT_Del52_ reduced its ability to bind Mpl (Figure 7E-G). Additional disruption of intermolecular ionic interactions involving the globular N-domain that are predicted to stabilize CRT_Del52_ dimers (Figure 5A) further decreases CRT_Del52_ binding to Mpl (Figures 7F and 7G) and eliminates its ability to induce proliferation of Ba/F3-Mpl cells in a cytokine-independent manner (Figures 7C and 7D). These results indicate an important role of charged residues at dimerization interfaces in stabilizing functionally active conformations of CRT_Del52._

The formation of Mpl_2_(CRT_Del52_)_2_ heterotetramers (~200 kDa) is previously suggested by size-exclusion chromatography (22). This could be achieved by binding of a (CRT_Del52_)_2_ dimer (Figure 5A) to two Mpl molecules, inducing Mpl dimerization (Figure 8). In this type of model, two interaction interfaces are predicted to be key determinants of mutant CRT-Mpl binding: (i) C-domain-dependent interactions conferring the Mpl targeting specificity to CRT mutants (consistent with Figure 2) and (ii) glycan-dependent interactions between N117 of Mpl and carbohydrate-binding site of the mutant CRT (consistent with published data (13, 22)). The loss of the majority or a subset of contacts with the first site are consistent with the impaired abilities of CRT_Del52Δ28_ and CRT_Del52Δ36_ to mediate cell proliferation (Figure 7A and (19)). Additionally, proliferation impairment observed with several dimer interface mutants (Figures 7C and 7D) is accompanied by altered disulfide/multimer patterns observed with those mutants (Figure 6).

**Figure 8:**
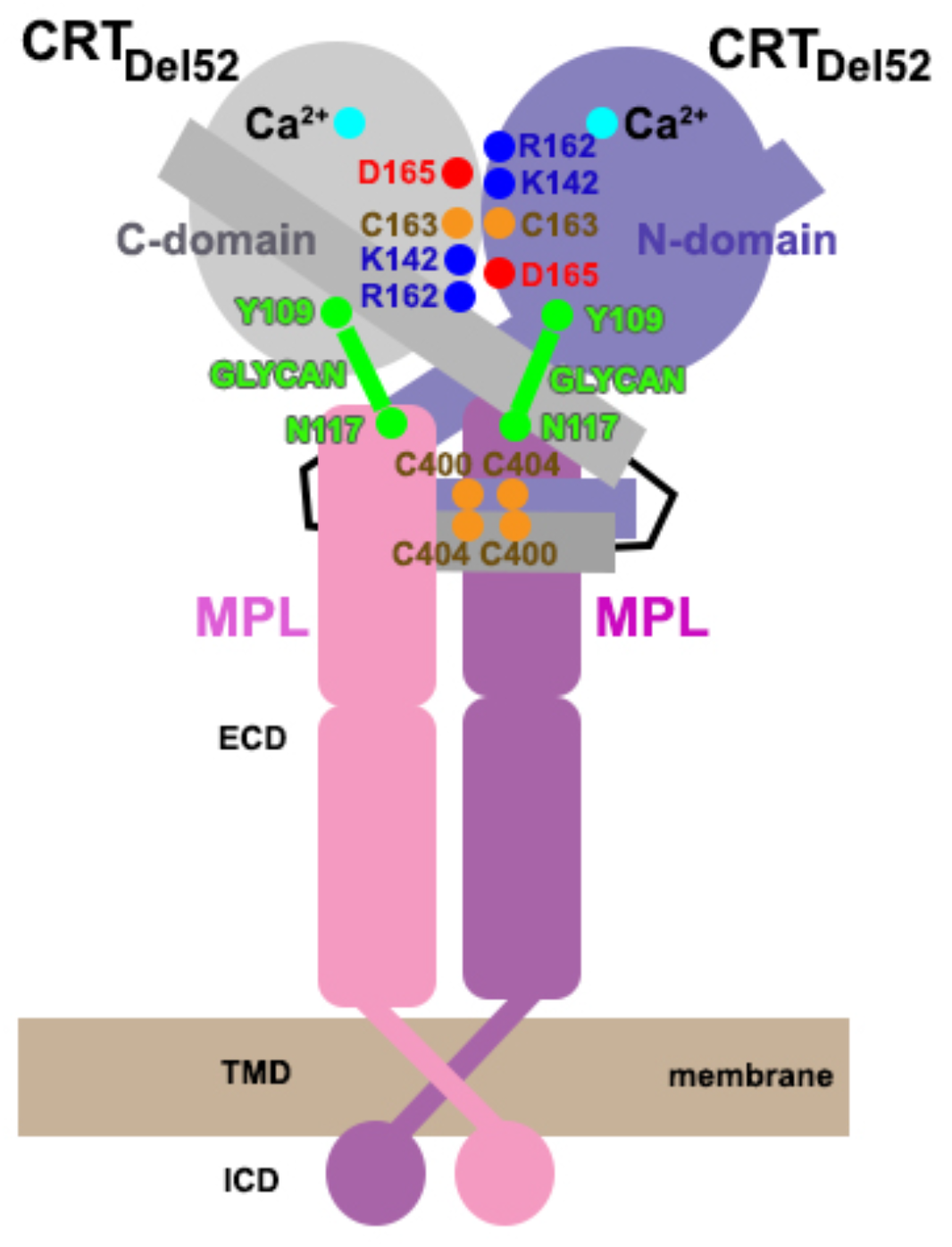
Schematic representation of a proposed Mpl_2_(CRT_Del52_)_2_ complex. Mpl activation is mediated by covalently-linked CRT_Del52_ dimers stabilized by disulfides and ionic interactions at C-domain and N-domain dimerization interfaces. The C-domain-of CRT_Del52_ also contributes to the specificity of Mpl binding (Figure 2), in addition to generic glycan-binding site residues of CRT that are previously shown to mediate Mpl recruitment (13, 20, 21).

Overall, the findings of this study reveal that disulfide-mediated CRT multimerization is a fundamental feature of MPN CRT mutants. The mutant sequences not only directly induce multimer formation via C-terminal disulfides, but also appear to induce a conformational change in the globular N-domain that promotes multimerization through that domain. It is noteworthy that C-terminal truncations of CRT from some species induce its oligomerization propensity via interactions which most likely resemble those induced within the MPN mutants (38). Thus, cancers-linked mutations of CRT confer selective growth advantages to cells by inducing specificity for Mpl, by driving multimerization of the receptor, and by exploiting a natural dimerization interface of CRT, that becomes relevant under cell-stress conditions.

## STAR Methods

### Healthy Donor and Patient Samples

Blood was collected after written informed consent in accordance with the University of Michigan Institutional Review Board approved protocols for a myeloproliferative diseases repository (HUM0006778) or the University of Michigan Platelet Pharmacology and Physiology Platelet core (HUM00107120).

### DNA Constructs

CRT_Ins5_ (K385fs*47 (3)) was made using CRT_WT_ (clone BC020493) as template by the QuickChange site-directed mutagenesis kit. CRT_Del52_ (L367fs*46 (3)) was amplified from a patient cDNA. All primers, vectors and cloning methods used for CRT and Mpl cloning are specified in the SI Materials and Methods. Point mutants were generated using the QuickChange site-directed mutagenesis kit.

### Antibodies

An anti-CRT mutant C-terminal antibody, anti-CRT(C_mut_), was generated by GeneTel Laboratories using rabbits immunized with a 22-mer peptide (KMSPARPRTSCREACLQGWTEA) derived from the CRT mutant C-terminus and affinity purified. Other antibodies are specified in the SI Materials and Methods.

### Platelet Isolation

Whole blood from healthy donors and MPN patients was collected in ACD anticoagulant tubes (BD Biosciences) and platelets isolated as described previously (39) with some modifications described in the SI Materials and Methods.

### Immunoblotting and Immunoprecipitations

Cell lines were lysed in lysis buffer (50 mM Tris pH 7.5, 150 mM NaCl, 1% Triton X-100, 5 mM CaCl_2_ and protease inhibitor cocktail (Roche)) for 1h at 4°C. For native, reducing and non-reducing lysate blot experiments, transfected HEK293T cells were lysed in the presence of 10 mM N-ethylmaleimide (Sigma). Protein concentration was measured using bicinchoninic acid assay (Pierce). For native blots, lysates were added to the loading buffer without reducing agents and SDS. For non-reducing blots, lysates were added to the loading buffer with SDS but without any reducing agents. For reducing blots, lysates were added to the loading buffer with SDS and reducing agent (65 mM DTT). Co-immunoprecipitation (IP) was performed using transfected HEK293T cells or Ba/F3 Mpl CRT cells as described (40) with some modifications described in the SI Materials and Methods.

### Cytokine independent proliferation

Ba/F3 Mpl cells expressing CRT constructs were seeded at 1×10^6^ cells/ml in the absence of mouse IL-3. Further, the proliferation rate of different cell lines was calculated by counting the live cells using a hemocytometer.

#### Data Sharing Statement

Data will be deposited in Dryad following manuscript acceptance.

### Statistical analysis

All statistical analysis was performed in GraphPad Prism (version 8.0c).

## Acknowledgements

We are grateful to all the blood donors for their contribution to this work. We thank the University of Michigan DNA Sequencing Core for DNA sequencing. We are grateful to Harihar Mohan, Angela Danielski and Maha Hamed for their contributions to the project. This work was funded by NIH grant (RO1 AI123957 to MR) and the University of Michigan Fast Forward Protein Folding Diseases Initiative. I.D.P. was supported by the NSF Division of Biological Infrastructure (award 1855425).

## Authorship Contributions

AV helped design experiments, performed experiments, helped analyze data, and wrote sections of the manuscript. JG helped design experiments, performed experiments related to Figures 3 and 7, helped analyze data and edited the manuscript. MK collected patient samples, purified platelets and edited the manuscript. IP generated the structural model for the CRT-Mpl tetrameric complex, wrote some sections, and edited the manuscript. MT directs the University of Michigan MPN repository and edited the manuscript. SW helped construct and sequence some CRT variants. AB helped construct and sequence some CRT variants described in this manuscript. MR designed the study, analyzed data, and wrote sections of the manuscript.

