## Supplemental Methods, Tables & Figures for "Mpl is activated by dimers of MPN-linked calreticulin mutants stabilized by disulfide bonds and ionic interactions"

**This file includes:**

Supplementary Methods

Tables S1 and S2

Figures S1 to S8

SI References

### SI Methods

#### Generation of CRT constructs

*Untagged $CRT_{WT}$, $CRT_{Ins5}$ and $CRT_{Del52}$ and their mutants constructs for mammalian expression*: Full-length untagged $CRT_{WT}$ and $CRT_{Ins5}$ were also cloned into the pMSCV-puro retroviral vector using XhoI and EcoRI sites. $CRT_{Del52}$ in the pMSCV vector was generated by replacing an AccI–EcoRI fragment of pMSCV-$CRT_{WT}$ with an AccI–EcoRI fragment of pGB1- $CRT_{Del52}$. Full-length $CRT_{WT}$, $CRT_{Ins5}$ and $CRT_{Del52}$ were sub-cloned into the pcDNA 3.1(-) zeo from the pMSCV puromycin vectors via restriction digestion using XhoI and EcoRI sites.

Cysteines mutants of full-length $CRT_{Ins5}$ (C163A, C419A, and C423A) and $CRT_{Del52}$ (C163A, C400A, C404A) were generated using the corresponding untagged pcDNA and pMSCV plasmids with the QuickChange site-directed mutagenesis kit. D165K, D166K, H170A mutations were also generated using the $CRT_{WT}$ or $CRT_{Del52}$ and $CRT_{Del52\text{-}3CA}$ ($CRT_{Del52(C163A/C400A/C404A)}$) untagged pcDNA and pMSCV plasmids using the QuickChange site-directed mutagenesis kit. C-terminal truncations of pMSCV-$CRT_{Del52}$ (Δ12, Δ19, Δ28 and Δ36) plasmids were generated by introducing stop codon at indicated positions using the QuickChange site-directed mutagenesis kit.

*GB1 and histidine-tagged C-domain constructs for mammalian expression*: The plasmid pcDNA 3.1/Zeo (-) from Invitrogen was used as the parent vector for construction of ligation independent cloning (LIC)-compatible mammalian expression vector (pcDNA-COX2-G) for expression of GB1 and histidine-tagged fusion proteins. The SspI restriction site in the plasmid was removed by site-directed mutagenesis and used as the parent vector for construction of the LIC-region-containing plasmids. The SspI-minus pcDNA 3.1 (-) vector was digested with XhoI and HindIII. The large DNA fragment from this digestion was gel purified. The LIC regions from pGB1 (DelProposto et al., 2009) (encoding $His_6$ and GB1 tag and TEV cleavage sites) were PCR amplified using the following oligos: 5'-GTCACTCGAGACCATGCACCATCATCATCATCATTCT-3', 5'GTCAAAGCTTGTCGACGGAGCTCGAATTCGGATC-3'. These oligos add a XhoI site followed by a modified Kozak sequence adjacent to the ATG translational start site of these regions and a HindIII site at the 3' end of the LIC region. These fragments were digested then ligated with the vector fragment. Positive clones were identified by PCR and confirmed by DNA sequencing. Using two rounds of PCR, pcDNA-COX2-G was constructed in which a secretion signal sequence was fused upstream of the polyhistidine sequence. The first round fused the signal sequence from Cyclooxygenase 2 (Cox2) to the LIC region, and the second round introduced the XhoI restriction site with the modified Kozak sequence and ATG to the end of the signal sequence. The following oligos were used to add the signal sequence to the end of the fusion LIC region: 5'GTGCGCAGTACTGGCTCTTTCTCACACTGCACACCATCATCATCATCAT and 5'CCTCTCGAGACCATGCTGGCTCGTGCACTGCTTCTGTGCGCAGTACTGGCT. The resulting fragments were digested with XhoI and HindIII, inserted into the vector and sequenced.

Using LIC, $CRT_{WT}$ (residues MKDKQDEEQRLKEEEEDKKRKEEEEAEDKEDDEDKDEDEEDEEDKEEDEEEDVPGQA KDEL), and $CRT_{Del52}$ C-domains (residues MKDKQDEEQRTRRMMRTKMRMRRMRRTRRKMRRKMSPARPRTSCREACLQGWTEA) were cloned into the pcDNA-COX2-G vector as N-terminal 6X histidine-tagged and GB1-tagged fusion proteins. Cysteine mutations (C44A, C48A) and truncations (Δ12, Δ19, Δ28 and Δ36) in the pcDNA-COX2-G-$CRT_{Del52}$ C-domain plasmids were also generated using the QuickChange site-directed mutagenesis kit.

*GB1 and histidine-tagged full-length $CRT_{WT}$, $CRT_{Del52}$ and $CRT_{Del52}$ truncation constructs for mammalian expression:* The full-length $CRT_{WT}$ was also cloned into pcDNA-COX2-G vector using LIC. $CRT_{Del52}$ in the pcDNA-COX2-G vector was generated by replacing the BsrGI–EcoRI fragment of pcDNA-COX2-G-$CRT_{WT}$ with the BsrGI–EcoRI fragment of pcDNA 3.1(-) $CRT_{Del52}$. C-terminal truncations of GB1 and histidine-tagged $CRT_{Del52}$ pcDNA-COX2-G (Δ12, Δ19, Δ28, and Δ36) plasmids were generated by introducing stop codon at indicated positions using the QuickChange site-directed mutagenesis kit.

*Mpl constructs*: cDNA of human Mpl (clone 731658 from DNASU) was cloned into the pMSCV-neo vector as untagged Mpl using EcoRI and XhoI sites or as an N-terminal $His_6$-FLAG fusion into pcDNA 3.1 (-) vector via LIC. pcDNA 3.1 (-) based-LIC plasmid encoding N-terminal $His_6$-FLAG fusion was generated in the same way as pcDNA-COX2-G except that the $His_6$-FLAG tag and TEV cleavage site was derived from pMCSG54 (http://bioinformatics.anl.gov/mcsg/technologies/vectors.html).

**Cell lines, Transient Transfections, and Infections**

Human embryonic kidney (HEK) 293T and phoenix cells were maintained in Dulbecco's Modified Eagle's Medium supplemented with 10% fetal calf serum and penicillin/streptomycin (100 U/ml, 100 mg/ml). Mouse proB cell line, Ba/F3 was maintained in RPMI 1640 supplemented with mouse IL-3 (BioLegend), 10% fetal calf serum and penicillin/streptomycin (100 U/ml, 100 mg/ml).

HEK293T cells were transfected with pcDNA 3.1(-) encoding indicated constructs using polyethylenimine (Polysciences, Inc) for 48 h. Retroviral supernatants encoding Mpl or CRT constructs were generated as previously described (Del Cid et al., 2010; Jeffery et al., 2011). Ba/F3 cells were first transduced with a retroviral vector encoding untagged human Mpl to generate Ba/F3-Mpl cells and selected with 1 mg/ml geneticin (Thermo Fisher) for Mpl expression. Ba/F3-Mpl cells were further transduced with retroviral vectors encoding various indicated CRTs or a control virus and subsequently selected using 1 mg/ml geneticin (Thermo Fisher) for Mpl expression and 3 μg/ml puromycin (Thermo Fisher) for CRT expression. For some experiments, full-length tagged CRT constructs in the pcDNA vector were nucleofected into Ba/F3-Mpl cells using cell line nucleofector kit V (Lonza) and selected using 0.2 mg/ml Zeocin.

**Platelet Isolation**

Platelet-rich plasma (PRP) was separated by centrifugation of whole blood at 200 x g for 15 minutes at room temperature with no brakes. PRP was extracted and treated with acid citrate dextrose (2.5% sodium citrate tribasic, 1.5% citric acid, 2.0% D-glucose) and apyrase (0.02 U/ml) (Sigma) and centrifuged for 10 min at 2000 x g. The platelet pellet was resuspended in Tyrode's buffer (10 mM HEPES pH 7.4, 134 mM NaCl, 12 mM $NaHCO_3$, 2.9 mM KCl, 0.34 mM $Na_2HPO_4$, 1 mM $MgCl_2$, 5 mM D-glucose) and rested at 37°C for 30 min before the use. The rested platelets in Tyrode's buffer were centrifuged at 2000 x g for 10 min and lysed in 500-800 μl of lysis buffer (50 mM Tris pH 7.5, 150 mM NaCl, 1% Triton X-100, 5 mM $CaCl_2$ and protease inhibitor cocktail (Roche)) using an end-over-end shaker for 30 min at 4°C.

**Protein Purification**

$CRT_{WT}$, $CRT_{Ins5}$ and $CRT_{Del52}$ in pGB1 vector were transformed into *Escherichia coli* Rosetta (DE3) cells. Bacterial cultures were induced with IPTG for 16-20 h at 20°C ($CRT_{WT}$) or 16°C

($CRT_{Ins5}$ and $CRT_{Del52}$). Proteins were purified by Nickel affinity chromatography as described previously (Del Cid et al., 2010). The GB1 and His tags were released using the Tobacco etch virus (TEV) protease cleavage as described (Wijeyesakere et al., 2011).

### Immunoblotting

Antibodies used in the immunoblotting analyses were: for $CRT_{WT,}$ a rabbit anti-CRT(N) antibody (Catalog number 12238; CST; 1:10000 dilution) or rabbit anti-CRT antibody (Catalog number PA3-900; Thermo-Fisher; 1:10000 dilution); for MPN CRT mutants, rabbit anti-CRT($C_{mut}$) antibody; for Mpl, rabbit anti-c-Mpl (Catalog number 06944; Millipore; 1:10000 dilution); for 6x His tag, mouse His-Tag antibody (Catalog number MA1-21315; Thermo-Fisher; 1:10000 dilution); for loading controls, anti-vinculin (Catalog number 13901; CST; 1:10000 dilution) and anti-GAPDH (Catalog number 5174; CST; 1:10000 dilution). Blots were incubated with a secondary antibody conjugated to horseradish peroxidase mouse anti-rabbit (Catalog number 211032171; Jackson Immunoresearch) or goat anti-mouse (Catalog number 115035003; Jackson Immunoresearch) and proteins were detected via chemiluminescence.

### Immunoprecipitations (IP)

Co-immunoprecipitation was performed using transfected HEK293T cells or Ba/F3 Mpl CRT cells as described (Del Cid et al., 2010) with some modifications. 500 μg of DTBP cross-linked cell lysates were incubated with custom anti-CRT($C_{mut}$) (2 μg) or anti-CRT (PA3-900; Thermo-Fisher; 1:1000) or anti-c-Mpl (06944; Millipore; 5 μg) antibodies over-night at 4°C. Similarly, healthy donor and MPN patient platelet lysates were lysed using the lysis buffer as described above but without cross-linking. Following clearance of the cell debris, 1 ml (0.5 mg) of the supernatants were incubated with either anti-CRT($C_{mut}$) (2 μg) or anti-c-Mpl (06944; Millipore; 5 μg) or without antibodies and mixed gently at 4°C for 16 h, centrifuged to remove cell debris, then incubated with protein G beads for 2 h, and beads washed with lysis buffer for 3 times and samples were eluted with Laemmli buffer. Samples were further denatured and separated by SDS-PAGE and immunoblotted as described above.

### Molecular modeling

Modeling and verification of functionally relevant $CRT_{Del52}$ dimers was performed in tow steps: (1) modeling of a monomeric $CRT_{Del52}$ (2) choice of dimerization mode from available crystal structures of CRT and (3) the verification of the functional relevance of the chosen $CRT_{Del52}$ dimer by mutational and functional studies.

We generated the monomer of $CRT_{Del52}$ using available atomic structure of human CRT (PDB ID: 5lk5 subunit E, for residues 19-203 and 303-366) (Moreau et al., 2016). The novel C-terminal domain was modeled as an α-helix extension, using fragment 367-386 from the structure of human PLC editing module (PDB ID: 6eny, subunit G) (Blees et al., 2017), with subsequent residue substitutions in accordance with the new C-terminal sequence of $CRT_{Del52}$ (Figure 2A). The distal C-terminal segment of $CRT_{Del52}$ ($^{399}$SCREACLQ$^{406}$) was modeled as a 2-turn α-helix, based on secondary structure predictions by I-TASSER (Roy et al., 2010), while the connecting loop (residues 387-398) was modeled in an extended conformation. Residue substitutions and modeling of an 8-residue C-terminal α-helical segment together with the 14-residue connecting loop were performed using PyMOL molecular graphic system (version 1.8.4.1 Schrodinger, LLC).

Second, we examined crystal structures of CRT oligomers (PDB IDs: 5klk5, 3pos, 3o0x) to identify dimerization modes that can account for the data described on Figures 1-4, consistent with formation of intermolecular disulfides between N-domains (C163-C163) and between C-domains (two C400-C404 disulfides) of monomers. Relatively stable dimers observed in crystal structure 5klk5 were used to build the models of $CRT_{Del52}$ dimers by molecular superposition. Two novel C-terminal α-helices of $CRT_{Del52}$ were brought together as dimers to form two disulfide bonds (C400-C404), shown to stabilize $CRT_{Del52}$ dimers (Figures 2 and 4).

Closer examination of the “N-N” dimerization mode of $CRT_{Del52}$ mutants (Figure 5A) allow predicting two additional symmetrical “N-C” dimerization interfaces that may be formed between N-domain of one molecule and C-domain of another molecule. A slight decrease of the α-helix kink at A352 would move the C-terminal part of the α-helix (residues 366-383) closer to N-domain glycan recognition site. This helix shift would bring positively charged and non-polar residues from C-domain of one molecule closer to negatively charged and aromatic residues from the N-domain glycan-binding site of the second molecule. As a result, dimer-stabilizing ionic and hydrophobic interactions could also be formed between the following residue pairs from N- and C-domains of interacting molecules: F74-M380, D125-R381, E127-R381, Y128-M377, Y109-T373, Y109-M377, Y109-R376, M131-T373, D135-R369, I147-M370, and D317-R376. The existence of ionic or hydrophobic interactions between these residue pairs may explain the significant role of residues 376-383 for multimerization of $CRT_{Del52}$ constructs, as their truncation in $CRT_{Del52\Delta36}$ eliminated oligomer formation (Figure 3).

To analyze the possible structures of higher order oligomers of $CRT_{Del52}$, we modeled tetramers of $CRT_{Del52}$ (Figure S6C). The modeling was performed by superposition of $CRT_{Del52}$ monomers with CRT trimer (subunits E, G, and J) from the crystal structure of 10-meric CRT complex for K71K mutant (PDB ID: 5lk5) (Moreau et al., 2016). To add the fourth $CRT_{Del52}$ subunit, we superposed it with an additional subunit X obtained by symmetry transformation of the crystal structure. In the resulting structure of a CRT tetramer, two subunit pairs are connected through C-terminal α-helices, while two other subunit pairs are connected through N-domain loops.

**Table S1.** List of MPN patient samples used in this study

| Patient ID | Age/Sex | Mutation | Platelets (K/μL) | Disease |
|---|---|---|---|---|
| 5502-1 | 44/F | CRT 52 bp del | 622 | ET |
| 5502-5 | 44/F | CRT 52 bp del | 441 | ET |
| 8744 | 75/M | CRT 4 bp del | 559 | ET |
| 1526 | 75/F | CRT 52 bp del | 270 | ET |
| 6105 | 31/F | CRT 52 bp del | 341 | ET |
| 2648-2 | 70/M | CRT 52 bp del | 389 | ET |
| 2648-3 | 70/M | CRT 52 bp del | 399 | ET |
| 6102 | 34/M | CRT 52 bp del | 312 | PMF |
| 4995-2 | 45/F | CRT 5 bp ins | 300 | ET |
| 3829 | 48/M | CRT 52 bp del | 738 | ET |
| 7245 | 30/F | CRT 52 bp del | 433 | ET |
| 8742 | 56/F | CRT 52 bp del | 178 | PMF |
| 8251-2 | 72/F | CRT 5 bp ins | 558 | PMF |
| 1718 | 81/F | CRT 52 bp del | 261 | PMF |
| 2791 | 62/M | CRT 52 bp del | 91 | post-ET MF |
| 1521 | 78/M | CRT 52 bp del | 273 | MF |
| 2028-1 | 66/M | CRT 52 bp del | 400 | post-ET MF |
| 1244 | 71/M | JAK2V617F | 98 | PMF |
| 9813 | 84/M | JAK2V617F | 420 | PMF |
| 2161 | 48/M | JAK2V617F | 422 | PV |
| 4493 | 69/F | JAK2V617F | 172 | post-PV MF |

PV, polycythemia vera; ET, essential thrombocythemia; PMF, primary myelofibrosis.

**Table S2**: List of primers used in this study.

| Target gene(s) | Primers | Purpose | Source or reference |
|---|---|---|---|
| $CRT_{Ins5}$ | Forward: 5'-GAGGAGGAGGAGGCAGAGGACAATTGTCGGAGGATGATGAGGACAAAG-3'; Reverse: 5'-CTTTGTCCTCATCATCCTCCGACAATTGTCCTCTGCCTCCTCCTCCTC-3' | QuickChange site directed mutation to create $CRT_{Ins5}$ | This study |
| $CRT_{WT}$ | Forward: 5'-AACTCGAGATGCTGCTATCCGTGCCGCTGCTGCTCGGC-3'; Reverse: 5'-AAGAATTCCTACAGCTCGTCCTTGGCCTGGCCGGGGACATCT-3' | Clone $CRT_{WT}$ into pMSCV puromycin | This study |
| $CRT_{Ins5}$ | Forward: 5'-AACTCGAGATGCTGCTATCCGTGCCGCTGCTGCTCGGC-3'; Reverse: 5'-AAGAATTCTCAGGCCTCAGTCCAGCCCTGGAGGCAG-3' | Clone $CRT_{Ins5}$ into pMSCV puromycin | This study |
| $CRT_{Del52}$, $CRT_{Ins5}$ | Forward: 5'-TACTTCCAATCCAATGCCGTCGCCGAGCCTGCCGTCTAC-3'; Reverse: 5'-TTATCCACTTCCAATGCTAGGCCTCAGTCCAGCCCTGGAG-3' | Clone $CRT_{Del52}$ and $CRT_{Ins5}$ into pGB1 | This study |
| CRT C-domain | Forward 5'-TACTTCCAATCCAATGCTATGAAGGACAAACAGGACGAG-3'; Reverse 5'-TTATCCACTTCCAATGCTAGGCCTCAGTCCAGCCCTGGAG-3' | Clone $CRT_{WT}$, $CRT_{Del52}$ and $CRT_{Ins5}$ C-domains into pcDNA COX-G using LIC | This study |
| $CRT_{Del52}$ (C400A/C404A); $CRT_{Ins5}$ (C419A/C423A); $CRT_{Del52\text{-}C}$ (C44A/C48A) C-domain | Forward 5'-CCAAGGACGAGCGCTAGAGAGGCCGCCCTCCAGGGCTG-3'; Reverse 5'-CAGCCCTGGAGGGCGGCCTCTCTAGCGCTCGTCCTTGG-3' | QuickChange site directed mutation to mutate novel cysteines | This study |
| $CRT_{Del52}$ (C163A) | Forward 5'-TGATCAACAAGGACATCCGTGCCAAGGATGATGAGTTTACAC-3'; Reverse 5'-GTGTAAACTCATCATCCTTGGCACGGATGTCCTTGTTGATCA-3' | QuickChange site directed mutation to mutate free cysteine C163 | This study |
| $CRT_{Del52}$ (D165K) | Forward 5'-ACAGGTGTGTAAACTCATCCTTCTTGCAACGGATGTCCTTG | QuickChange site directed mutation to mutate D165 | This study |

| | | | |
|---|---|---|---|
| | -3'; Reverse 5'-CAAGGACATCCGTTGCAAGAAGGATGAGTTTACACACCTGT-3' | | |
| $CRT_{Del52}$ (D166K) | Forward 5'-GTACAGGTGTGTAAACTCCTTATCCTTGCAACGGATGTCC-3'; Reverse 5'-GGACATCCGTTGCAAGGATAAGGAGTTTACACACCTGTAC -3' | QuickChange site directed mutation to mutate D166 | This study |
| $CRT_{Del52}$ (D165K/D166K) | Forward 5'-CAGTGTGTACAGGTGTGTAAACTCCTTCTTCTTGCAACGGATGTCCTTGTTGAT-3'; Reverse 5'-ATCAACAAGGACATCCGTTGCAAGAAGAAGGAGTTTACACACCTGTACACACTG -3' | QuickChange site directed mutation to mutate D165 and D166 | This study |
| $CRT_{Del52}$ (C163A/C400A/C404A/D165K) | Forward 5'-ACAGGTGTGTAAACTCATCCTTCTTGGCACGGATGTCCTTG-3'; Reverse 5'-CAAGGACATCCGTGCCAAGAAGGATGAGTTTACACACCTGT-3' | QuickChange site directed mutation to mutate D165 in $CRT_{Del52}$ 3CA | This study |
| $CRT_{Del52}$ (C163A/C400A/C404A/D166K) | Forward 5'-GTACAGGTGTGTAAACTCCTTATCCTTGGCACGGATGTCC-3'; Reverse 5'-GGACATCCGTGCCAAGGATAAGGAGTTTACACACCTGTAC-3' | QuickChange site directed mutation to mutate D166 in $CRT_{Del52}$ 3CA | This study |
| $CRT_{Del52}$ (C163A/C400A/C404A/D165K/D166K) | Forward 5'-GTGTGTACAGGTGTGTAAACTCCTTCTTCTTGGCACGGATGTCCTTGTTG-3'; Reverse 5'-CAACAAGGACATCCGTGCCAAGAAGAAGGAGTTTACACACCTGTACACAC-3' | QuickChange site directed mutation to mutate D165 and D166 in $CRT_{Del52}$ 3CA | This study |
| $CRT_{Del52\Delta12}$ | Forward 5'-GGCAGGCCTCTCTCTAGCTCGTCCTTGGCC-3'; Reverse 5'-GGCCAAGGACGAGCTAGAGAGAGGCCTGCC -3' | QuickChange site directed mutation to create Δ12 truncation | This study |
| $CRT_{Del52\Delta19}$ | Forward 5'-GTCCTTGGCCTGGCCTAGGACATCTTCCTCCT-3'; Reverse 5'-AGGAGGAAGATGTCCTAGGCCAGGCCAAGGAC-3' | QuickChange site directed mutation to create Δ19 truncation | This study |
| $CRT_{Del52\Delta28}$ | Forward 5'-CCTCATCTTCCTCTATGTCCTCCTCATCCTCCTCATC-3'; Reverse 5'-GATGAGGAGGATGAGGAGGACATAGAGGAAGATGAGG-3' | QuickChange site directed mutation to create Δ28 truncation | This study |

| | | | |
|---|---|---|---|
| $CRT_{Del52\Delta36}$ | Forward 5'-CCTCCTCATCCTCCTCATCTACATCTTTGTCCTCATCAT-3'; Reverse 5'-ATGATGAGGACAAAGATGTAGATGAGGAGGATGAGGAGG-3' | QuickChange site directed mutation to create Δ36 truncation | This study |
| Mpl | Forward 5'-TACTTCCAATCCAATGCCCAAGATGTCTCCTTGCTGG-3'; Reverse 5'-TTATCCACTTCCAATGCTAAGGCTGCTGCCAATAGCTTAG-3' | Clone Mpl into pcDNA COX2-54 | This study |
| Mpl | Forward 5'-CCCGAATTCATGCCCTCCTGGGCCCTCTTC-3', reverse 5'-CCCCTCGAGTCAAGGCTGCTGCCAATAGCTT-3' | Clone Mpl into pMSCV neomycin | This study |

**Figures and legends**

**SI Figures**

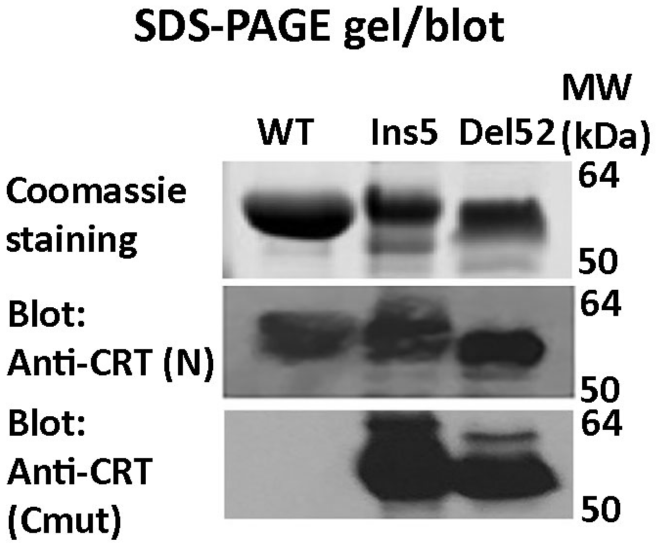


**Fig. S1: Verification of the specificity of anti-CRT($C_{mut}$) antibody**
$CRT_{WT}$, $CRT_{Ins5}$ and $CRT_{Del52}$ were purified following expression in *E. coli*. Concentration matched (20 µg) TEV-digested $CRT_{WT}$, $CRT_{Ins5}$ and $CRT_{Del52}$ proteins were separated by SDS-PAGE and Coomassie stained (top panel) or immunoblotted with anti-CRT(N) (middle panel) or anti-CRT($C_{mut}$) (bottom panel) antibodies.

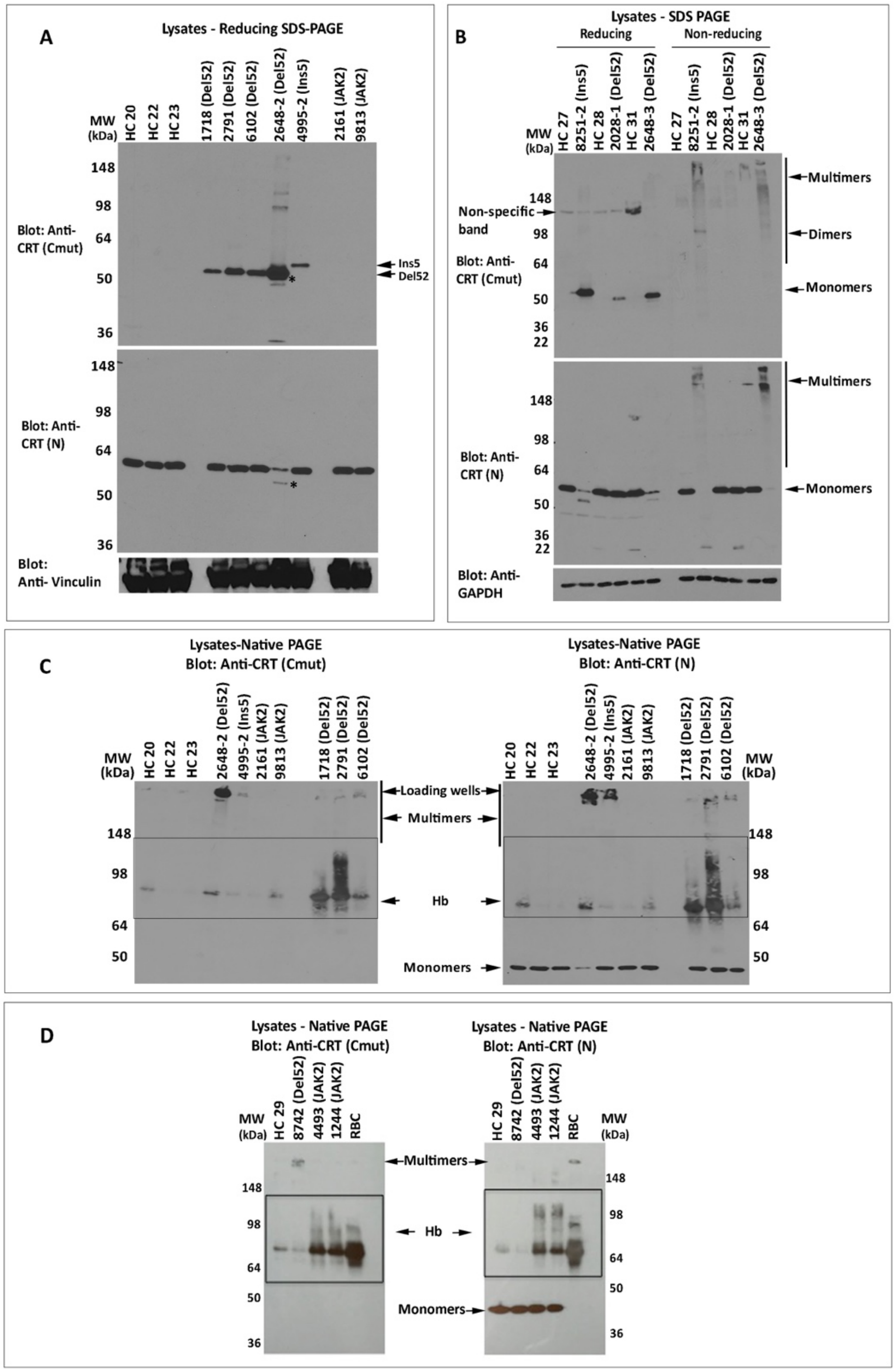


**Fig. S2: Mutant CRT is multimeric via disulfide bonds in MPN patient platelets**

A) SDS-PAGE (8% gels) under reducing conditions and immunoblots of platelet lysates from healthy donors and MPN patient platelets were probed with the indicated antibodies. The same lysates were loaded onto two gels, and probed with anti-CRT($C_{mut}$) (top panel) or anti-CRT(N) (middle panel) antibodies. The latter was re-probed with anti-vinculin (lower panel) antibody. Anti-CRT($C_{mut}$) does not detect wild type CRT in healthy control lysates. Lysate 4995 is a $CRT_{Ins5}$ mutant, which based on size (Figure 2A), should migrate more slowly than $CRT_{Del52}$ (all other samples in the blot). B) Lysates from MPN patients or healthy donor platelets were separated by reducing or non-reducing SDS-PAGE (4-20% gradient gels) and the same lysates were loaded onto two gels probed with anti-CRT($C_{mut}$) (top panel) or anti-CRT(N) (middle panel) antibodies. The former was re-probed with anti-GAPDH antibody. Monomer and multimer CRT bands are indicated. Bands indicated as dimers or multimers are over-represented under non-reducing conditions in the CRT mutant platelet lysates, whereas monomer bands are depleted under the same conditions for 8251-2 and 2648-3, which appear to have a high mutational load. The lower expression of mutant CRT in 2028-1 patient lysate precluded its detection under non-reducing conditions, at greater than background levels. Anti-CRT(Cmut) does not detect wild type CRT in healthy control lysates, hence no bands are visualized in lanes labeled HC in immunoblots with anti-CRT($C_{mut}$). Non-specific bands that are detected in the reducing blots are marked as such. C-D) Native immunoblots (8% gels) of platelet lysates (25 μg) from MPN patient platelets or same-day healthy donor platelets or red blood cell lysates, were probed with anti-CRT($C_{mut}$) (C, D left panels) and anti-CRT(N) (C, D right panels) antibodies. Oligomeric forms of mutant CRT are readily detectable at the top of the gel, particularly with samples such as 2648-2, which showed high expression of mutant CRT. Boxes indicate Hb contamination of platelets, verified using red blood cell (RBC) lysates. CRT HC indicates healthy control samples; CRT mutant patient samples are indicated as Del52 or Ins5; JAK2 indicates JAK2 mutant patient samples. RBC indicates red blood cell lysates.

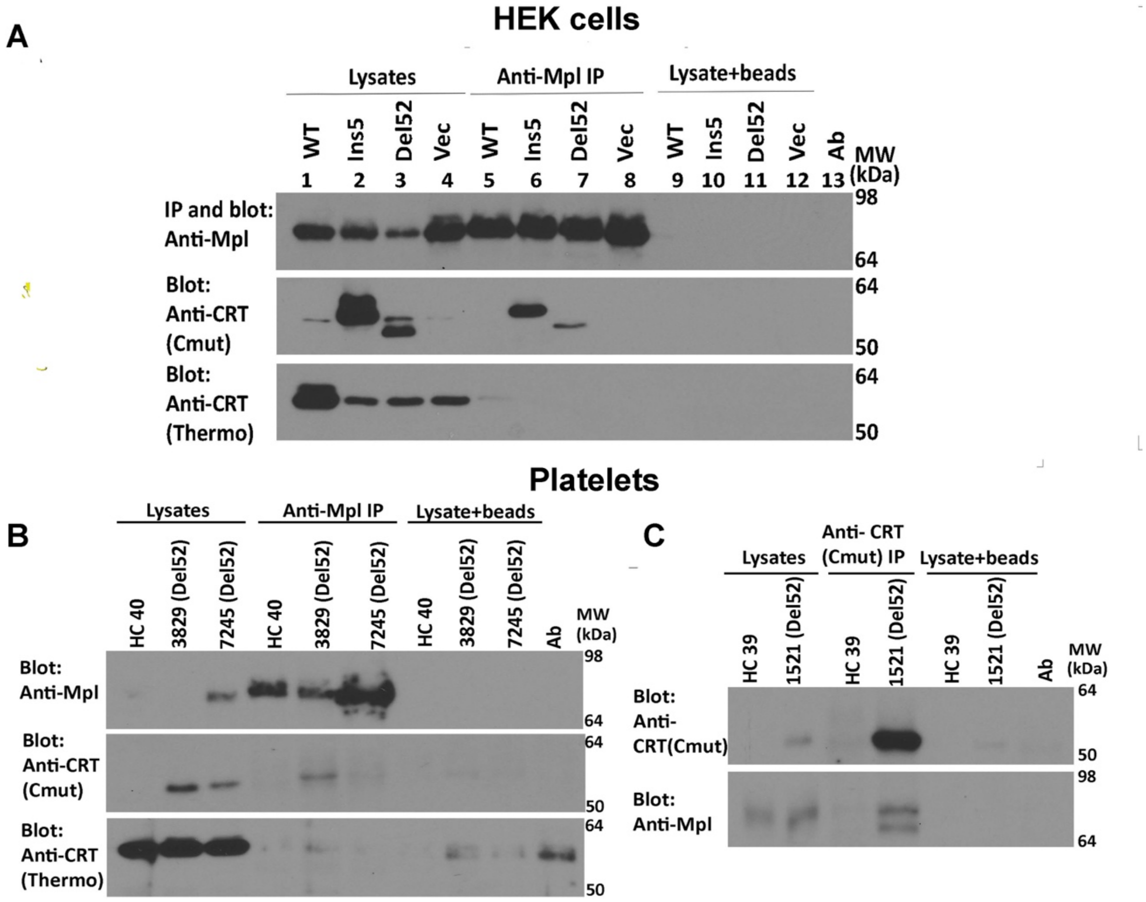


**Fig. S3: Mutant CRT preferentially interact with Mpl in HEK293T cells and MPN patient platelets**

A) Lysates from HEK293T cells expressing full-length untagged wild type or mutant CRTs (Ins5 or Del52) and Mpl, or control cells expressing Mpl alone (Vec) were immunoprecipitated (indicated as IP) with anti-Mpl antibody and subsequent immunoblotting was undertaken with the indicated antibodies. Data shown are representative of 3 independent experiments. B and C) Lysates from indicated healthy donor (HC) or MPN patient platelets (Del52) were immunoprecipitated with anti-Mpl antibody (B) or anti-CRT($C_{mut}$) antibody (C) and subsequently analyzed by immunoblotting with the indicated antibodies. HC represents healthy donor; CRT mutant patients are indicated as Del52. In all panels, non-specific interactions in the absence of primary antibody are shown by the lysate+ beads lanes.

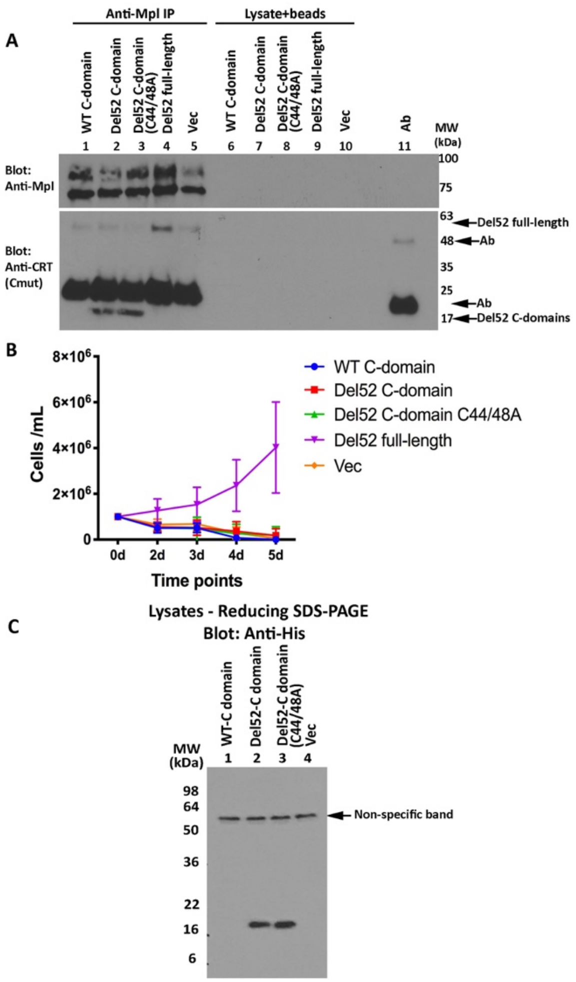


**Fig. S4: The isolated C-domain of $CRT_{Del52}$ is expressed at higher levels than its wild type counterpart and binds Mpl, but is not sufficient for inducing cytokine-independent cell proliferation**

A) Lysates from Ba/F3-Mpl cells expressing N-terminal His and GB1-tagged C-domains of $CRT_{WT}$, $CRT_{Del52}$, $CRT_{Del52(C44/48A)}$, or untagged full-length $CRT_{Del52}$ or control cells expressing untagged Mpl alone (Vec) were immunoprecipitated (indicated as IP) with anti-Mpl antibody and subsequent immunoblotting was undertaken with the indicated antibodies. The full-length $CRT_{Del52}$, $CRT_{Del52}$ C-domain and antibody (Ab)-derived bands are indicated. Results are representative of 4 independent experiments from two separate transductions of Ba/F3-Mpl cells with the indicated viruses. Non-specific precipitation in the absence of primary antibody is shown by the lysate+beads lanes. B) Cytokine-independent proliferation by $CRT_{Del52}$ C-domain mutants. Ba/F3-Mpl cells were transduced with retroviral vectors encoding N-terminal His and GB1-tagged C-domains (sequences indicated in Figure 2A) of $CRT_{WT}$, $CRT_{Del52}$, $CRT_{Del52(C44A/C48A),}$ or full-length untagged $CRT_{Del52}$ or a control virus and subsequently cultured in the absence of mouse IL3 and proliferation was measured based on cell counting on the indicated days. Data are averaged from two separate transductions of Ba/F3-Mpl cells with the indicated viruses, of a total of 4 analyses. C) Expression levels of $CRT_{Del52}$ C-domains are higher than WT C-domain even in the absence of Mpl. Lysates from HEK cells expressing N-terminal His and GB1-tagged C-domains of $CRT_{WT}$, $CRT_{Del52}$, $CRT_{Del52(C44/48A)}$ were separated by SDS-PAGE (4-20% gradient gel) and analyzed by immunoblotting with anti-His antibody. The expression of the WT C-domain is not detectable under conditions where mutant C-domains are readily detectable.

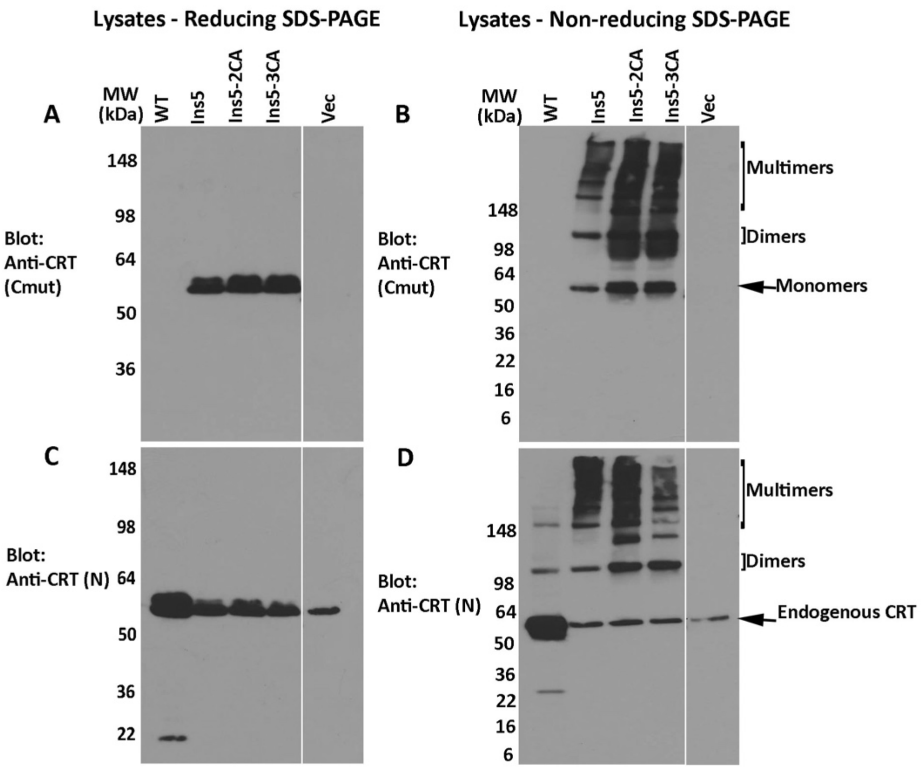


**Fig. S5: $CRT_{Ins5}$ mutant forms disulfide-linked multimers**

HEK293T cells were transiently transfected with plasmids encoding full-length untagged $CRT_{WT}$, $CRT_{Ins5}$, $CRT_{Ins5\text{-}2CA}$ ($CRT_{Ins5(C419A/C423A)}$), $CRT_{Ins5\text{-}3CA}$ ($CRT_{Ins5(C163A/C419A/C423A)}$), or plasmid lacking CRT (Vec). Cell lysates from indicated cells were separated by SDS-PAGE under reducing (8% gels) (A, C) or non-reducing (4-20% gradient gels) (B, D) conditions and immunoblotted with indicated antibodies. Data are representative of 3 sets of analyses. In blots following non-reducing SDS-PAGE (B and D), species consistent with the size of CRT monomers, dimers and endogenous CRT are indicated.

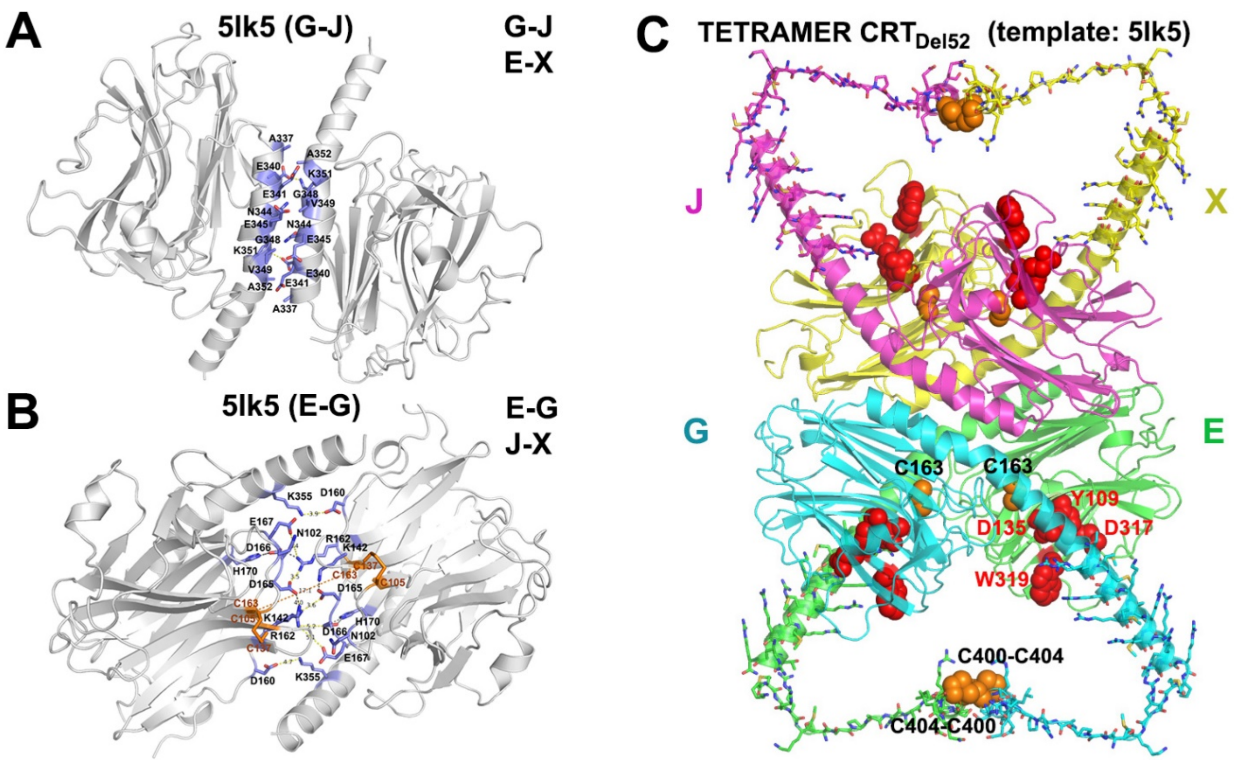


**Fig. S6. Modeling of CRT$_{Del52}$ tetramer using C-domain helix-helix and N-domain loop-loop dimerization modes.**

Two major dimerization modes were observed in the crystal structure of the 10-mer complex of CRT D71K mutant (PDB ID: 5lk5 (Moreau et al., 2016)): (A) tight packing of antiparallel α-helices ("C-C" dimer between subunits G-J and E-X) and (B) dimerization via N-domain loops rich on charged residues that form intermolecular ionic bridges ("N-N" dimer, between subunits E-G and J-X). Contacting residues at C-domain helix-helix and N-domain loop-loop interfaces are shown by sticks colored blue for C-atoms. (C) A homology model of CRT$_{Del52}$ tetramer lacking P-domains was generated using both types of dimers observed in the crystal structure of CRT D71K mutant. The molecular model is shown by a cartoon colored green, cyan, purple, yellow for subunits E, G, J, X, respectively. Subunit X absent in the structure of 10-mer was obtained by symmetry transformation. Mutated residues (367-406) from the novel C-tail of CRT$_{Del52}$ are shown by sticks. Cysteine residues from each subunit predicted to form intermolecular disulfide bonds are shown by orange spheres. Residues (Y109, D135, D317, W319) located in glycan-binding pockets of globular domains (Kozlov et al., 2010) are shown by red spheres.

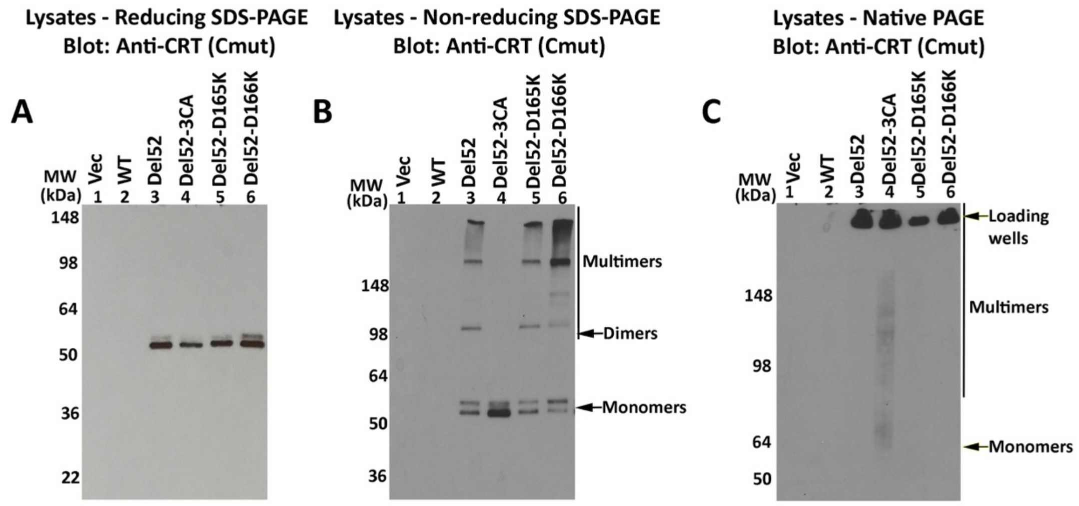


**Fig. S7: Multimer formation by Del52-D165K and Del52-D166K mutants**

HEK293T cells were transiently transfected with plasmids encoding untagged full-length $CRT_{Del52}$, $CRT_{Del52\text{-}D165K}$, $CRT_{Del52\text{-}D166K}$ or $CRT_{Del52\text{-}3CA}$ ($CRT_{Del52(C163A/C400A/C404A)}$) or a control vector. Cell lysates from indicated cells were separated by SDS-PAGE under reducing (8% gels) (A) or non-reducing (4-20% gradient gels) (B) conditions or by native-PAGE (8% gels) (C) and immunoblotted with anti-CRT($C_{mut}$) antibody. In blots following SDS-PAGE under non-reducing and native conditions, species consistent with the size of mutant CRT monomers, dimers, multimers are indicated. Location of loading wells is also indicated.

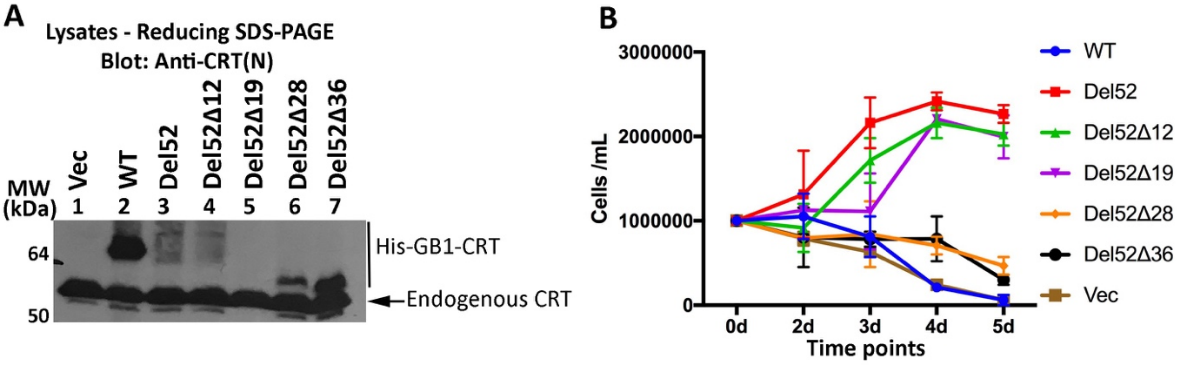


**Fig. S8: Cytokine-independent proliferation induced by truncated His-GB1-tagged $CRT_{Del52}$ mutants.**

Ba/F3-Mpl cells were electroporated with pcDNA-CoxG plasmids encoding N-terminal His-GB1 tagged full-length $CRT_{WT}$, $CRT_{Del52}$, $CRT_{Del52\Delta12,}$ $CRT_{Del52\Delta19,}$ $CRT_{Del52\Delta28,}$ $CRT_{Del52\Delta36}$ or a control vector and subsequently selected with zeocin at 0.2 mg/ml. A) Lysates from Ba/F3-Mpl cells expressing the indicated constructs were immunoblotted with anti-CRT(N) antibody (single analysis). B) Cells were subsequently cultured in the absence of mouse IL3 and proliferation was measured based on cell counting on the indicated days. Data are averaged from 2 independent proliferation experiments undertaken following a single electroporation of Ba/F3-Mpl cells.
